# Top-down perceptual inference shaping the activity of early visual cortex

**DOI:** 10.1101/2023.11.29.569262

**Authors:** Ferenc Csikor, Balázs Meszéna, Katalin Ócsai, Gergő Orbán

## Abstract

Deep discriminative models provide remarkable insights into hierarchical processing in the brain by predicting neural activity along the visual pathway. However, these models differ from biological systems in their computational and architectural properties. Unlike biological systems, they require teaching signals for supervised learning. Moreover, they rely on feed-forward processing of stimuli, which contrasts with the extensive top-down connections in the ventral pathway. Here, we address both issues by developing a hierarchical deep generative model and show that it predicts an extensive set of experimental results in the primary and secondary visual cortices (V1 and V2). We show that the widely documented sensitivity of V2 neurons to textures is a consequence of learning a hierarchical representation of natural images. Further, we show that top-down influences are inherent to hierarchical inference. Hierarchical inference explains neural signatures of top-down interactions and reveals how higher-level representation shapes low-level representations through modulation of response mean and noise correlations in V1.

---

Hierarchical processing of visual information is a fundamental property of the visual cortex. Recently, deep learning models have emerged as central tools to investigate the hierarchical processing of visual information [1]. These models rely on a feed-forward processing hierarchy, which enables them to effectively process natural images. A highly successful class of models applied to the visual system are goal-oriented image models, which postulate that the relevant training objective of visual cortical computations is to perform specific tasks, such as classification of inputs into discrete categories. Such goal-directed models have been immensely successful in predicting neuronal responses to natural images, such that progression of processing stages qualitatively matched those of the ventral stream [2], as well as in designing synthetic images that elicit specific response patterns in populations of visual cortical neurons [3, 4]. However, images that were specifically designed to confuse the feed-forward model also significantly delayed neural responses, indicating that computations beyond feed-forward processing were recruited by the visual system [5].

Top-down interactions between the processing stages are ubiquitous in the visual cortex (Fig. 1) but lack a well-established role in goal-directed models of vision. Scrutinizing the computational principles underlying the goal directed models might provide normative arguments why biological vision recruits a more complex architecture and how top-down connections help perceptual processes. Goal-oriented models learn to compress stimulus information such that information relevant for the specific task is retained. For instance, if a model is trained for animal classification, it will excel in this task by recognizing a tiger irrespective of its posture. To achieve this, goal-directed models rely on a learning paradigm called supervised learning, which capitalizes on image and category pairs to train the hierarchical model. Recently studies have actually highlighted that the specific task the goal-oriented model is trained on is actually affecting how well a model accounts for neural data [6]. In contrast with supervised learning of a task-specific neural representation, biological learning imposes two seemingly contradicting requirements. First, representations need to be learned without supervision, i.e., learning in the visual cortex should be performed without a signal what its output should be. Second, the visual system is expected to learn a model of natural images that is not adapted to perform one specific task, instead one that can flexibly perform arbitrary tasks, according to the needs of the current context of the animal. In other words, while under some circumstances it is sufficient to infer if we are facing a tiger or a zebra, in other circumstances it is imperative to precisely judge which movements are plausible based on the current posture of the animal. However, optimization for only one set of task variables (distinguishing different species) does not provide guarantees for efficient computations relying on other variables (evaluating the posture). These requirements motivate a task-independent framework for hierarchical processing in the visual cortex [7].

**Fig. 1.**
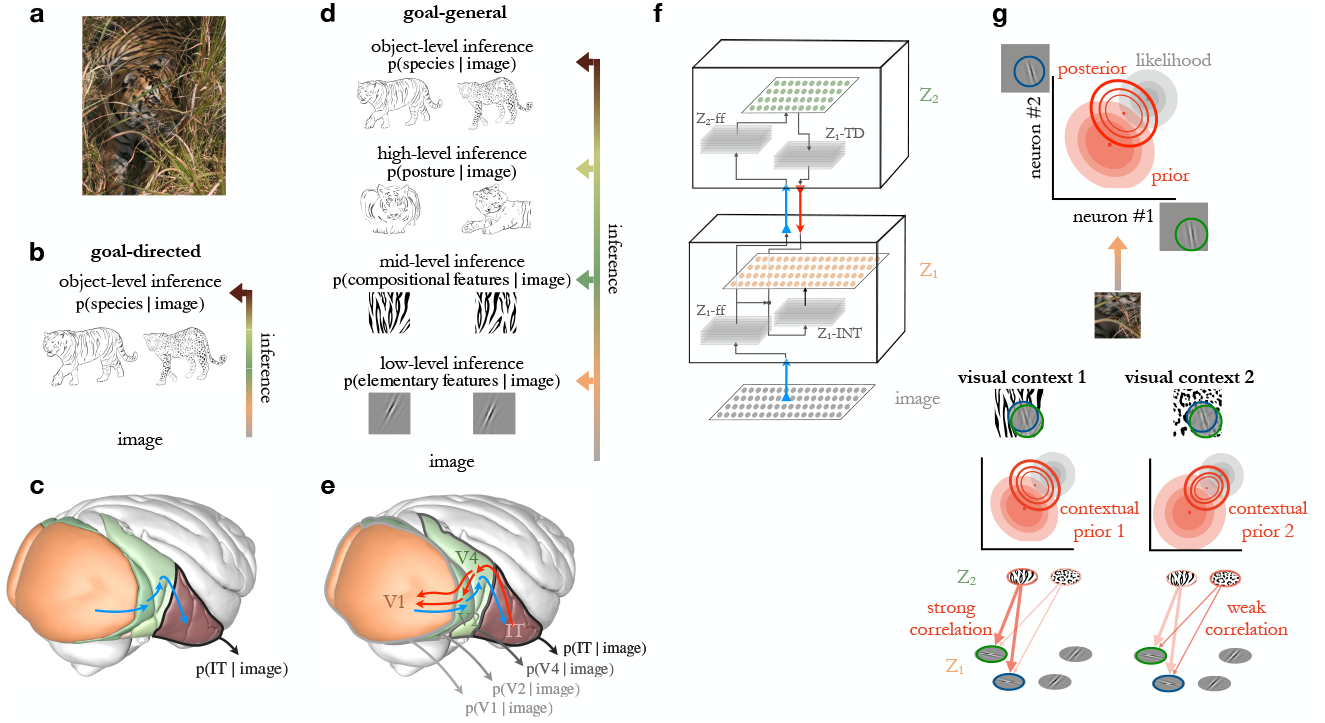
Hierarchical inference in task-independent models. **a**, Example natural image. **b**, Inference in a goal-directed model, which aims to identify categories of images. **c**, Illustration of feed-forward processing in the ventral stream of the visual cortex. **d**, Inference in a hierarchical task-independent model, which permits reverse engineering the contribution of a hierarchical set of features to the observed stimulus. **e**, Illustration of top-down (*red arrows*) influences supplementing feed-forward processing (*blue arrows*) along the hierarchy of the ventral stream [10]. **f**, Neuronal circuitry of TDVAE that learns a hierarchy of features Z_1_ (*orange*) and Z_2_ (*green*) corresponding to V1 and V2 in the ventral stream. Features are represented in a layer of neurons (*planes with orange and green disks* that correspond to individual neurons). Layers of neurons (*gray planes*) transform feed-forward (Z_1_-ff, Z_2_-ff, *blue arrows*) and top-down information (Z_1_-TD, *red arrow*) to ensure precise inference. Integration of feed-forward and top-down information is achieved at Z_1_-INT. **g**, Computational role of top-down connections. *Top*: Response intensities of a pair of Z_1_ neurons (axes) are determined by the features they represent (insets on axes). Interpretation of the image (posterior distribution, *empty contours*) is the combination of the prior (*filled red ellipses*) and the evidence carried by the image (likelihood, *gray ellipses*). Mean response of the neuron (measured across trials, or across a long span of time) is the posterior mean (*dot*), its correlation is contributing to the noise correlation. *Middle*: Change of contextual priors upon changes in stimulus statistics. *Bottom*: Contribution of Z_2_ neurons to contextual priors in Z_1_. *Saturated colors* indicate stronger activation. Image credit (a): istockphoto/Getty images.

We propose that the ingredient that can address the dual challenge of learning is provided by top-down interactions. We argue that top-down signals have a central role in a class of task-independent learning systems, which has recently achieved considerable success in machine learning applications, deep generative models. Conceptually, top-down interactions establish expectations to support inference [8]: representing and computing with expectations is a key topic in theories of perception [9, 10] and was shown to have both neuronal and behavioral correlates [8, 11, 12]. Top-down interactions in the hierarchy ensure that instead of a simple feed-forward pass of information to perform inference at the top of the processing hierarchy, inferences in intermediate layers of processing are supported by contextual information fed back from higher levels of the hierarchy. Formally, top-down interactions establish a contextual prior which is used to integrate with the feed-forward information channel to perform hierarchical inference.

In this study we investigate how a task-independent hierarchical model is acquired through adaptation to natural image statistics. Specifically, we investigate the hierarchical representations learned in a deep generative model from two major directions. First, we seek to characterize learned representations to see if it reflects the properties of the early and mid-level visual cortices, V1 and V2 in particular. Second, we seek to understand if top-down effects found in V1 are aligned with the way high-level representations interact with low-level representations by establishing contextual priors. In order to explore the defining characteristics of this model, we study texture representations, which is motivated by three factors. First, textures are highly relevant features of natural images and critically contribute to higher-level visual processes such as segmentation and object recognition [13, 14]. Second, texture representations have been established in mid-level visual cortices, and in particular in V2, across multiple species, ranging from humans [15], through non-human primates [16, 17, 18], to rodents [19], suggesting that textures are not only relevant from a machine learning point of view but from a neuroscience point of view as well. Third, while some features that the V2 is sensitive to can be identified with linear computations, textures can only be learned through non-linear computations, a critical component of the successes of deep learning models.

Based on fundamental principles of hierarchical inference, we extend the feed-forward deep learning framework to perform task-independent hierarchical inference (Fig. 1). The adopted Variational Autoencoder (VAE) framework allows unsupervised learning of a nonlinear model of natural images. We develop the hierarchical TDVAE (Top-Down VAE) model, which we train end-to-end on natural image patches. First, to establish the feasibility of the model we contrast the representation learned by the hierarchical TDVAE with the responses of neurons in the early visual cortex of macaques. Next, we investigate the learned properties of top-down connections through a series of experiments that explore two aspects: contextual influences of V2 on V1 responses, and effects of V2 representations on V1 response properties. Through the development of a Variational Autoencoder model, our work provides a normative interpretation of the extensive top-down network present in the visual cortical hierarchy and demonstrates signatures of top-down interactions both in the mean and correlations of neural activities.

## Results

To investigate the contribution of top-down connections to computations in a normative framework, we introduce a task-independent hierarchical model of natural images. We introduce our task-independent framework by contrasting it with alternative frameworks. The objective of hierarchical computations can be phrased in two different ways. First, the objective can be to learn to perform inference about the presence of high-level visual features. The most common form of this objective is a goal-directed objective, in which high-level features are object categories occurring in natural images (Fig. 1b). Here, *p*(object | image) is calculated such that object category information is provided during the training to learn a flexible and non-linear mapping from images to object categories [1, 2, 20]. A related objective in which high-level features are inferred is a self-supervised paradigm in which the goal is to identify features through learning invariances in images [21, 22]. In both of these models, intermediate sets of features are learned, with increasingly more complex features identified in the processing pipeline, which serve the inference of the features at the top level (in the case of goal-directed models, object categories). Here, neurons in the ventral stream of the visual cortex are identified with the different stages of a feed-forward pipeline (Fig. 1c). Second, a more general objective can be established by requiring the learning machinery to infer not only the presence of objects but also the presence of lower-level features constituting objects (Fig. 1d). This is phrased as learning a joint distribution over objects and features of different complexities: *p*(features_L_, features_M_, features_H_, object | image). This is a more fine-grained form of inference that enables performing not only one particular task, such as viewpoint invariant recognition of objects, but also learning to infer the different aspects of different observations of the same visual component. Given that a task-independent learning is taking place, no category labels are provided at any level of the computational hierarchy. To establish a connection between the model and the activity of cortical neurons, we interpret the activations of features in the model as the activation of model neurons. We identify the activity of model neurons with that of neurons in the ventral stream of the visual cortex, such that features at different levels of the model hierarchy correspond to neurons in different cortical regions (Fig. 1e).

To gain insight into the computations necessary for task-independent hierarchical inference, we focus on inference on two levels of features: *p*(features_L_, features_M_ | image), which we will refer to as Z_1_ and Z_2_ for clarity. Thus, the values in vector Z_1_ denote the activity of a population of low-level model neurons, while values in Z_2_ denote the activities of model neurons at the higher layer of the computational hierarchy. Image-evoked activity in Z_1_ is formulated as an inference problem:

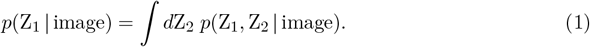

Here, *p*(Z_1_ | image), the posterior distribution, assigns probabilities to different response intensities of the neurons of the Z_1_ population upon observing a stimulus such that the most likely response intensity corresponds to the most probable value of the feature in the image. Probability theory helps to simplify the calculation of response distributions in individual areas. It defines two possible partitioning of the joint distribution *p*(Z_1_, Z_2_ | image). The *feedforward* partitioning solely relies on feed-forward progression of information (see also Methods): *p*(Z_1_ |image)·*p*(Z_2_ |Z_1_). However, the *top-down* partitioning also relies on top-down interactions between processing stages: *p*(Z_1_ |Z_2_, image)·*p*(Z_2_ | image). Note that in the latter formulation the activity in Z_1_ not only depends on the stimulus, but also on the activity in Z_2_, a signature of top-down influences. These formulations provide mathematically equivalent formulations to the inference problem and as such either of these could be used to learn a hierarchical task-independent model. However, as inference in such models is intractable, approximations are needed. Interestingly, the equivalence of feed-forward and top-down partitioning holds for exact inference but some of the most effective approximations, variational inference, terminates equivalence [23]. This, together with the top-down interactions characteristic of the anatomy of the visual cortex [24] motivates us to focus on hierarchical inference that recruits top-down interactions besides feed-forward processing of stimuli (Fig. 1e). To control for a possible effect of this choice, we implement the feed-forward parametrization as well. In our study we will seek to establish links between model neurons in Z_1_ and Z_2_ with neurons in V1 and V2 regions of the visual cortex.

In order to train a network to perform hierarchical inference, we turn to the Variational Autoencoder formalism [25, 26]. Variational Autoencoders (VAEs) have been developed to learn nonlinear task-independent image models. VAEs learn a pair of models: the recognition model supports inference, i.e. it describes the probability that a particular feature combination underlies an observed image; and the generative model, which summarizes the ‘mechanics’ of the environment, i.e. how the combination of features produces observations (see Methods for details). Relying on advances in hierarchical versions of VAEs [27, 28], we developed Top-Down VAE, or TDVAE for short (Fig. 1f). When constructing the TDVAE model, we parameterized a VAE relying on two principles. First, we capitalized on neuroscience intuitions about V1 and V2 to construct the architecture for learning about low- and mid-level features, Z_1_ and Z_2_. Second, the architecture is determined by a highly principled approximation in probabilistic inference but apart from the well-motivated restrictions (optimizing a lower bound to the true objective, performing variational amortized inference, deterministic propagation of signals across layers of the hierarchy), the TDVAE was left as general as possible such that its properties were determined solely by natural image statistics. Along this line, we assumed an energy-efficient representation of the natural environment in the generative model [29], which encouraged sparseness in population responses (Methods). Further, we assumed a linear relationship between the features that Z_1_ neurons are sensitive to and the pixels of natural images [29, 30]. At the level of Z_2_, the generative model allowed flexible nonlinear computations such that more complex computations and more complex feature sets can be learned. Next, we relied on anatomical insights when implementing inference: the top-down formulation *p*(Z_1_ |Z_2_, image)·*p*(Z_2_ | image) identifies two components: *p*(Z_1_ |Z_2_, image), and *p*(Z_2_| image). These components are implemented by neural networks. To implement these, we incorporated the anatomical constraint that information does not bypass V1 before reaching V2. This was achieved by sharing the entry layers of the two networks, later forking into a feed-forward neural network converging onto Z_2_, and a feedback neural network that integrated the information fed back from Z_2_ (see also Methods). Such choices are widely used in the machine learning literature and are referred to as parameter sharing. In order to demonstrate that the shared feed-forward pathway does not limit the performance of the TDVAE model, we have also implemented the alternative model with distinct feed-forward pathways. In summary, computations in TDVAE are visualized in a computational graph. This describes the way the input is processed through subsequent layers of processing to infer Z_1_ through feedback from Z_2_ and how the generative component contributes to the reconstruction of the input, the precision of which ultimately drives learning (Extended Data Fig. 1). In our paper we focus on the analysis of the recognition model, that is, activities of model neurons in Z_1_ and Z_2_ are used to predict patterns in cortical neural activities (Fig. 1f). The resulting circuitry of the recognition model was thus determined by the principles of probability theory and the task-independent training objective, termed Evidence Lower Bound, ELBO (Fig. 1f, see Methods for details).

The top-down component in TDVAE provides a key insight about the role of top-down connections: When the information carried by observations is limited by poor viewing conditions, occlusion, or noise, the interpretation of an input image bears the burden of uncertainty (Fig. 1a). Uncertainty implies that an observer needs to rely on their prior to interpret the scene [31, 32] (Fig. 1g, top), which contains information about learned regularities in the environment. Importantly, regularities change from environment to environment, implying that an observer knowledgeable about the environment shall rely on local environment-dependent priors, termed contextual priors (Fig. 1g, middle). Thus, when interpreting an image, Z_2_ neurons integrate information from a wide field of view (due to large receptive fields), and provide contextual priors for Z_1_ neurons through top-down connection patterns that reflect the statistics of the local context. In different environments different activation patterns in Z_2_ alter the contextual information of Z_1_ neurons (Fig. 1g, bottom).

Since the visual cortex was adapted to the natural environment [31, 32], the TDVAE model was trained end-to-end on natural image patches (Methods), such that the learned representations (i.e. the represented sets of features) are determined by the natural image statistics. The primary determinants of the learned model are the properties of the generative model while the recognition model was set to be sufficiently complex to accommodate the computations required for inference. To demonstrate the invariance of our results on the specific choice of the generative and recognition models, we trained an array of models with different parametrizations, including the feed-forward partitioning of the posterior (Eq. 1) and an untrained TDVAE model (see Methods). The goodness of these alternative models was established through calculating the

ELBO (Fig. 2a, Extended Data Table 4, Methods) and the one having the highest ELBO was used in the main text of the study, while alternatives are presented as Extended Data. During the construction of the architecture we argued that anatomical intuitions suggest a departure from the most general form of VAE as TDVAE shares the parameters of the circuitries serving Z_1_ and Z_2_ inference. To ensure that this choice does not affect model performance, we contrasted the shared feed-forward pathway model with the distinct pathway model. Simulations did not identify major performance setbacks with constraining the model to shared pathways (ELBOs are −400 and −407 for the 20-pixel shared and distinct pathway models, respectively, see also Methods). Apart from these architectural choices, TDVAE has no free parameters that are fitted to neural data, all parameters are determined by natural image statistics.

**Fig. 2.**
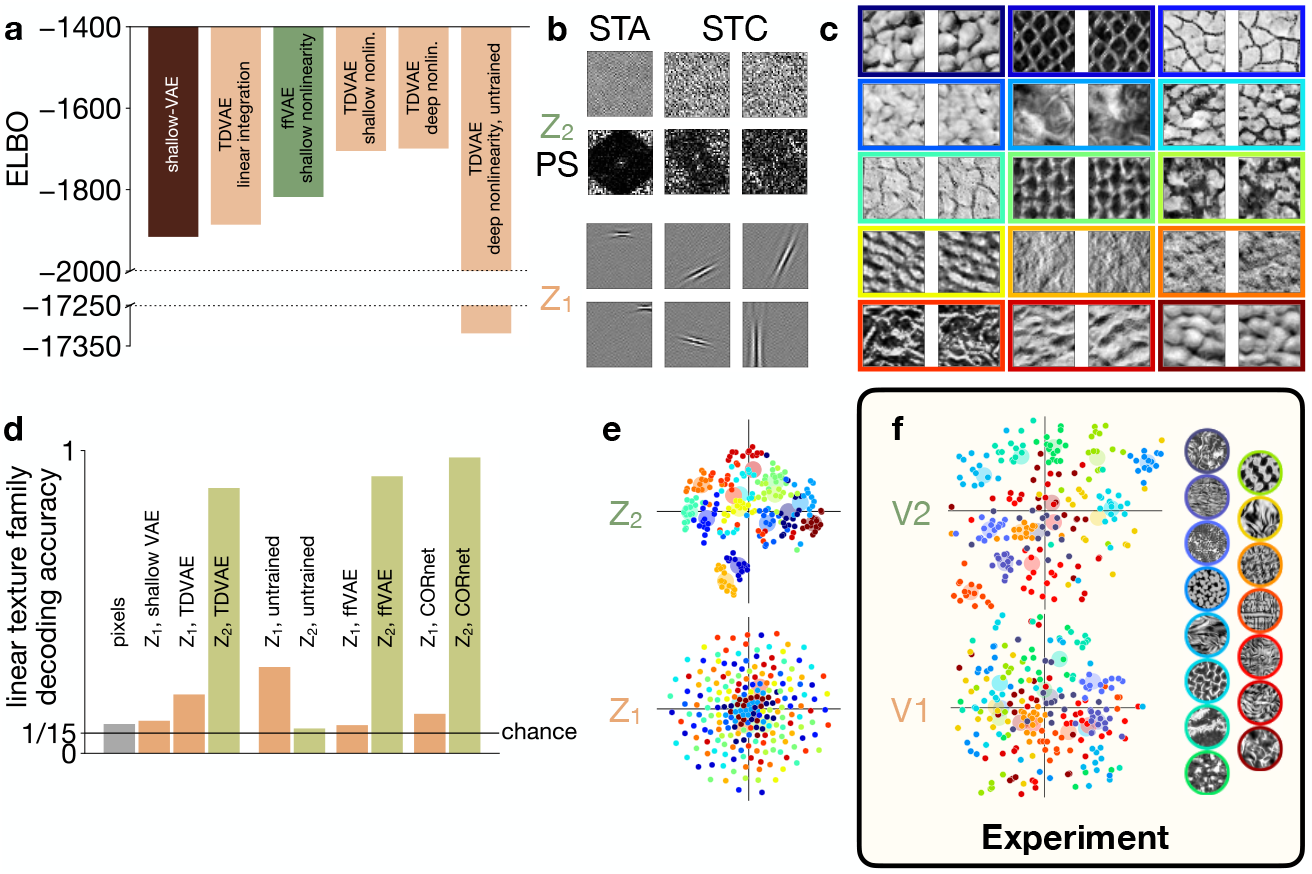
Hierarchical representation in Z_1_ and Z_2_ learned by TDVAE. **a**, Loss function of alternative models. *Shallow-VAE*: non-hierarchical VAE constrained to Z_1_; *ffVAE*: hierarchical VAE with feed-forward processing; *TDVAE*: variants of hierarchical VAEs with top-down processing. *Linear integration*: feed-forward and top-down pathways are linearly combined to reach Z_1_; *shallow nonlinearity*: generative model between Z_2_ and Z_1_ relies on single-layer MLP (Extended Data Table 3); *deep nonlinearity*: as above but with two-layer MLP. *Untrained*: Model with random parameters. **b**, *Top*: The first-order, and two example second-order receptive fields (STA and STC, respectively, see Methods) of a selected active Z_2_ neuron, together with power spectra (PS). *Bottom*: Example first-order receptive fields of six Z_1_ units. **c**, Examples of texture families (colors). **d**, Texture family decoding accuracies from different layers of the hierarchy. Means of *n* = 5 fits are shown; s.d. *<* 0.01 everywhere. *Black line*: chance performance. *Pixels*: direct decoding from images. Labels as in **a**. *CORnet*: goal-directed feed-forward model. **e**, Two-dimensional visualization (t-SNE) of mean responses of Z_1_ and Z_2_ neurons to randomly sampled texture images (*dots*). *Colors* as on panel c. *Disks*: mean across samples from a family. **f**, Same as **e** but in V1 and V2 recordings from macaques. Reproduced with permission from [17](PNAS).

Activation of features at the level of Z_1_ and Z_2_ in TDVAE are assumed to correspond to activations of V1 and V2 neurons up to a linear combination of the variables (see Methods for details), standard in earlier approaches [2]. The probabilistic nature of inference (Eq. 1) necessitates that we define how probability distributions are represented. A deterministic approach focuses on the best interpretation of the image, thus the neural response would be identified with the maximum of the distribution. In neural data, this maximum a posteriori response is identified with the trial-averaged response and variance is interpreted as mere noise in the circuitry. In contrast, optimal computations require a probabilistic interpretation. For this, we adopt the sampling hypothesis [33]. According to the sampling hypothesis, neural activity at any given time is a stochastic sample taken from the probability distribution and the sequence of samples constitutes the distribution, with mean of the neural activity corresponding to the most probable interpretation of the stimulus, and variance of the responses corresponding to the uncertainty associated with the interpretation (see also Methods). Thus, mean of *p*(Z_1_ |image) and *p*(Z_2_ |image), referred to as posteriors, corresponds to the mean response of the V1 and V2 neurons. Spike count collected in a time window corresponds to a finite number of samples from the posterior therefore it is expected to show considerable variance around the mean posterior, which can be reduced through averaging across trials. At the level of V1, a single sample can be identified with an activity from a 20 ms time window [34]. Width of the posteriors is reflected in response variance, while correlations in the posterior are reflected in noise correlations. Importantly, noise correlations can be produced by various sources [35], and thus our interpretation corresponds to a form of noise correlation that results from high-level perceptual variables. In this paper response variances are not investigated in detail but we refer the reader to earlier work [36, 37], and later sections discuss response correlations. Activity of model neurons are not constrained to be positive and therefore are identified with membrane potentials, similar to earlier approaches [36] (see Methods for details). The presented results can be translated to firing rates by transforming membrane potentials with a threshold-linear transformation [36, 38]. We decided to report results on the untransformed activities of Z_1_ and Z_2_ in order to demonstrate that the results of an end-to-end natural image trained VAE can be directly applied to predict neural data.

### Hierarchical representation of natural images

First, we investigate the representations that emerge in TDVAE. At the level of Z_1_, we found that TDVAE robustly learns a complete dictionary of Gabor-like filters (Fig. 2b, Extended Data Fig. 6, Extended Data Table 4). The receptive fields of individual model neurons are localized, oriented, and bandpass (Extended Data Fig. 7), qualitatively matching those obtained by training a VAE lacking a Z_2_ layer (Extended Data Fig. 8). This result confirms earlier studies using single layer linear generative models featuring sparsity constraint on neural activations [29, 30]. Thus, learning a hierarchical representation complete with a Z_2_ layer on top of Z_1_ left the qualitative features of the Z_1_ representation intact in the TDVAE model.

Unlike the compact receptive fields of Z_1_ model neurons, the representation at the Z_2_ layer has no noticeable linear structure (Fig. 2b STA). The second-order, nonlinear receptive fields of Z_2_ units, however, reveal orientation and wavelength selectivity (Fig. 2b STC, Methods), offering a first glimpse into the representation learned by the Z_2_ layer.

Inspired by extensive evidence supporting that V2 neurons are sensitive to texture-like structures [15, 19, 39], we used synthetic texture patches to further explore the properties of the Z_2_ representation that TDVAE learned on natural images [40]. We selected 15 natural textures characterized by dominant wavelengths compatible with the image patch size used in the study (Fig. 2c, Methods). We matched low-level texture family statistics (Methods), thus making texture families linearly indistinguishable in the pixel space (Fig. 2d, Methods). Since model Z_1_ neurons learn a linear mapping from the neural space to the pixel space, we expect that texture information cannot be read out with a linear decoder from Z_1_ response intensities either. Decoders that queried the texture family that a particular image was sampled from but ignored the identity of the image showed low performance on model Z_1_ neurons, which was still distinct from chance (0.1943 ± 0.0012, error denotes standard deviation over five repeats of the fit, Fig. 2d). The non-hierarchical version of VAE that completely lacked a Z_2_ was slightly outperformed by TDVAE (0.1073 ± 0.0020, Fig. 2d).

Nonlinear components in the generative model endow the Z_2_ to build a representation that is capable of distinguishing texture families such that textures can be effectively represented in this layer. Texture was reliably decodable from the mean responses of Z_2_ neurons of TDVAE (0.8761 ± 0.0001, Fig. 2d). This texture representation was very robust as it was qualitatively similar for all investigated hierarchical architectures (Extended Data Table 5). The texture representation discovered by TDVAE was low dimensional, irrespective of the architectural details of the model (Extended Data Table 5). This finding is surprising but recent experiments have confirmed that even though textures are represented by large neuron populations in the V2 of macaques, the effective dimensionality of this representation is very limited [19]. Intriguingly, the TDVAE architecture best fitting natural images only featured texture-selective dimensions, yielding a very compact representation exclusively focusing on textures (Extended Data Tables 4, 5). Responses of the Z_2_ population displayed clustering by the texture family (Fig. 2e), similar to the representations identified in V2 neuron populations in macaque (Fig. 2f). In contrast, no such clustering could be identified in Z_1_ (Fig. 2e), reminiscent of V1 neuron populations of the same macaque recordings (Fig. 2f).

We introduced three alternative models as controls to our TDVAE model. First, we investigated a model that had identical architecture to the TDVAE but was not adapted to natural images. Second, we tested the robustness of the learned texture representation against the form of partitioning of the posterior distribution. Third, representation learned by a goal directed feed-forward model was tested. The untrained model showed slightly higher decoding performance on texture families in Z_1_ (0.2850 ± 0.0078) but, in contrast with TDVAE, markedly lower performance in Z_2_ than the TDVAE (0.0820 ± 0.0076, Fig. 2d). The contra-intuitive higher Z_1_ result is explained by the fact that decoding is performed from a large dimensional space (matching the dimensionality of active Z_1_ dimensions) and the non-linear recognition model has the potential to distribute responses of Z_1_ such that it carries texture family information. As it is the generative model that assumes linear relationship between Z_1_ and pixel activations, the nonlinear recognition model ‘unlearns’ texture decodability as the generative and recognition models are jointly adapted to natural images. In contrast with TDVAE, no clustering was found in either Z_1_ or Z_2_ of the untrained model (Extended Data Fig. 9a).

The variant of TDVAE using feed-forward partitioning of the joint posterior distribution instead of top-down partitioning, ffVAE (*feed-forward VAE*), showed limited texture family decodability from Z_1_ (0.0930 ± 0.0003) and high texture family decodability from Z_2_ (0.9150 ± 0.0002, Fig. 2d), similar to TDVAE. This underlines the robustness of the learned texture representation against the exact implementation of the recognition model.

We have contrasted the hierarchical representation learned by TDVAE with that of a goaldirected model. We used a standard feed-forward implementation available in the literature, the CORnet model [41]. The CORnet model was trained on ImageNet and we designed analyses analogous to those performed with TDVAE (see Methods). In particular, we sought to identify layers across the processing hierarchy that bear similarities with Z_1_ and Z_2_ of TDVAE. Early layers with localized and orientation sensitive filters could be identified in this goal-directed model. Texture family decoding was not possible from the responses of this layer (0.1303 ± 0.0005), but layers immediately above this layer were characterized by a representation from which texture family decoding was effective (0.9770 ± 0.0003, Fig. 2d).

We investigated the representation learned by TDVAE further in order to contrast it with more specific properties of the representation found in V1 and V2. We first investigated the learned invariances of TDVAE. Mean responses of individual Z_1_ neurons displayed a high level of variability both across instances of texture images belonging to the same texture family and those belonging to different texture families (Fig. 3a). This tendency was similar to the sensitivities displayed by V1 neurons in the macaque visual cortex (Fig. 3b). In contrast, Z_2_ neurons displayed a higher level of invariance across texture images belonging to the same texture family (Fig. 3a), again reflecting the properties of macaque V2 neurons (Fig. 3b). To quantify this observation, across-population statistics was calculated by dissecting across-sample and across-family variances in mean responses both in Z_1_ and Z_2_ neurons of TDVAE using a nested ANOVA method [17]. Note, that in order to make our analysis more general by permitting an arbitrary linear mapping between model Z_2_ neurons and biological neurons, we used a random rotation of Z_2_ to perform the analysis (Methods). The relative magnitude of response variance across families versus across samples was significantly higher in Z_2_ than in Z_1_ (Fig. 3c, geometric means of variance ratios: Z_1_: 0.064, Z_2_: 2.097; one-sided independent two-sample t-test in the log domain, P = 4.3 × 10^−82^, t = −32.7, df = 198), corroborating experimental findings (Fig. 3d).

**Fig. 3.**
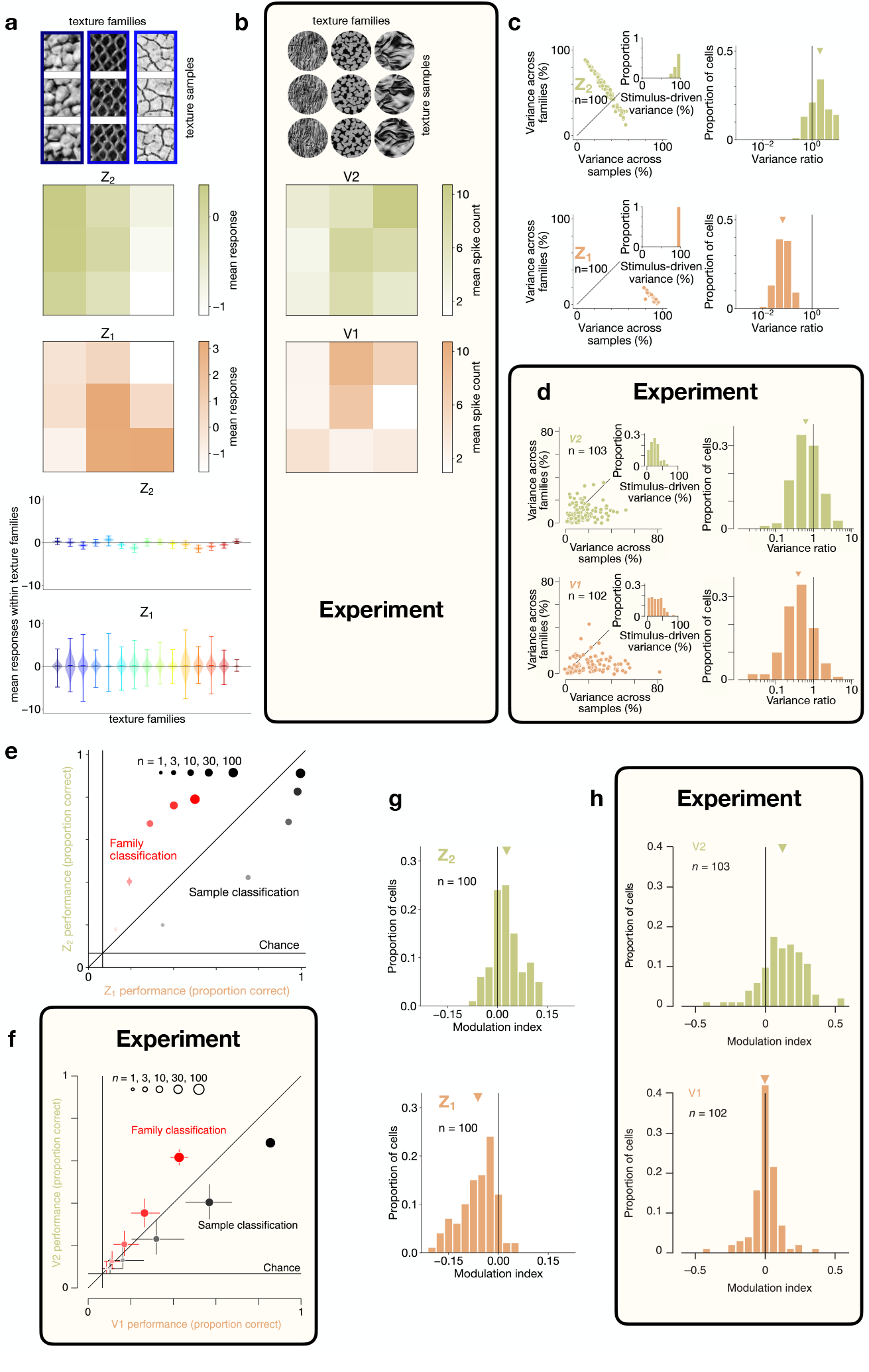
Signatures of learning a texture representation. **a**, Mean responses of example single neurons from Z_2_ and Z_1_ (rows two and three, respectively) to three example images from three texture families (*top*). *Bottom*: Variability of mean responses across 4000 images within, and across texture families (center lines: means, whiskers: extrema). **b**, Same as **a** in V1 and V2 recordings from macaque monkeys. Reproduced with permission from [17](PNAS). **c**, *Left*: Partitioning single-unit response variances into variance across images from different families, variance across images within families and a residual component (across stimulus repetitions) with nested ANOVA both in Z_2_ and Z_1_ (*n* = 100 neurons, *dots*). *Insets*: Distribution of the sum of the first two components of variance. *Right*: Distribution of the ratio of the first two components (*triangles* indicate geometric means). **d**, Same as **c** in 102 V1 and 103 V2 units from a pool of 13 macaque monkeys. **e**, Decoding texture family (*red*) and within-family stimulus identity (*black*) from the same 100 Z_1_ and Z_2_ units (*horizontal axis* and *vertical axis*, respectively) as in **c.** *Dot sizes*: decoding on randomly selected subpopulations (*n* = 1, 3, 10, 30, 100). *Solid line*: Chance performance. Dot centers: means; error bars: 95% confidence intervals of bootstrapping over included neurons and partitionings. **f**, Same as **e** from single unit V1 and V2 macaque recordings. **g**, Phase scrambling-induced modulation of the magnitude of mean responses to texture images in the same 100 Z_2_ and Z_1_ neurons (*top* and *bottom*, respectively) as in **c. h**, Same as **g** from V1 and V2 recordings in macaques. Experimental data is reproduced with permission from [16] and [17] (Nature Neuroscience, reproduced with permission from SNCSC, and PNAS, respectively).

Higher invariance of responses to different stimuli belonging to the same texture family in Z_2_ than in Z_1_ can also be captured by a linear decoding analysis (Fig. 3e). Efficiency of discrimination between images in the same texture family is weaker in Z_2_ than that of Z_1_ (Fig. 3e). Such increasing invariance to lower-level features along the visual hierarchy is a hallmark of gradual compression and can be identified at the population level in the primate visual cortex as well (Fig. 3f). In contrast with within-family stimulus identity decoding, decoding of texture family is more efficient from Z_2_ than from Z_1_ in TDVAE (Fig. 3e), a finding confirmed by macaque recordings (Fig. 3f).

Texture families can be defined through a set of pairwise statistics over the co-activations of linear filters represented in Z_1_/V1 [40]. To demonstrate that the learned Z_2_ representation reflects these quintessential features of a proper texture representation, the TDVAE model can be tested with stimuli in which these high-level statistics are selectively manipulated while keeping the low-level statistics intact (Methods). To achieve this, we used phase scrambling of images [16]. As texture sensitive neurons are assumed to be sensitive to high-level statistics, their removal is expected to reduce Z_2_ activations. For this, a modulation index was calculated for each neuron, which was defined as the difference between sample- and presentation-averaged absolute responses to undisturbed texture images and their phase scrambled counterparts normalized by their sum. Texture-sensitive units in Z_2_ showed higher modulation than active units in Z_1_ (Fig. 3g, mean modulation indices: Z_1_: −0.062, Z_2_: 0.028; one-sided independent twosample t-test, P = 2.6 × 10^−27^, t = −12.6, df = 198), in line with findings in macaque recordings (Fig. 3h).

In summary, Z_1_ and Z_2_ representations of TDVAE display key features of V1 and V2 representations of the macaque visual cortex. In particular, an elaborate texture representation is a salient feature of hierarchical representations of natural images.

### Top-down contributions to hierarchical inference

In a task-independent model inference can be performed on all levels of the hierarchy (Fig. 1d). According to equation (1), top-down influences affect the inference of Z_1_ activities through establishing a contextual prior for Z_1_ (Fig. 1g). Importantly, contrary to simple Bayesian computations, this prior is not an invariant component, instead it depends on the higher-level interpretation of the scene. In the following sections we seek to identify signatures of this contextual prior in Z_1_ response statistics: first on mean responses, second on response correlations.

Top-down influences can be expected when local interpretation of images depends on the surroundings. Establishing a wider context is directly supported by the growing receptive field sizes in higher hierarchical layers of TDVAE. To study top-down influences in detail, we rely on an implementation of TDVAE, which is trained on larger (50-pixel, see Methods) natural image patches than our standard model, which was trained on 40-pixel images. While the larger patch size requires more careful training, it offers a more precise evaluation of contextual effects. A particularly strong form of contextual effect can be established by selectively blocking direct stimulus effects from designated Z_1_ neurons and therefore top-down effects can be studied in isolation by investigating the emerging Z_1_ activation. This ‘illusory’ activity in Z_1_ neurons resonates with the interpretation of illusions as the contribution of priors to perception under uncertainty [42]. We constructed illusory contour stimuli by creating Kanizsa square stimuli (Fig. 4a) such that illusory contour segments were aligned to individual Z_1_ neurons and measured mean responses (Fig. 4b). We compared Kanizsa-elicited responses to both unobstructed images of squares and to control stimuli that were built from identical elements to the Kanizsa stimuli but in configurations not congruent with the percept of a square. Illusory edge responses were similar to unobstructed square stimulus-evoked responses, albeit with a lower gain for the illusory edge (Fig. 4b), similar to responses of V1 neurons of macaques [43] (Fig. 4c). Confirming expectations, responses to Kanizsa-incongruent stimulus elements were muted (Fig. 4b), again reproducing experimental findings (Fig. 4c). This tendency was consistent across the model neurons investigated using stimuli that were tailored to their response properties (see Methods, Fig. 4d).

**Fig. 4.**
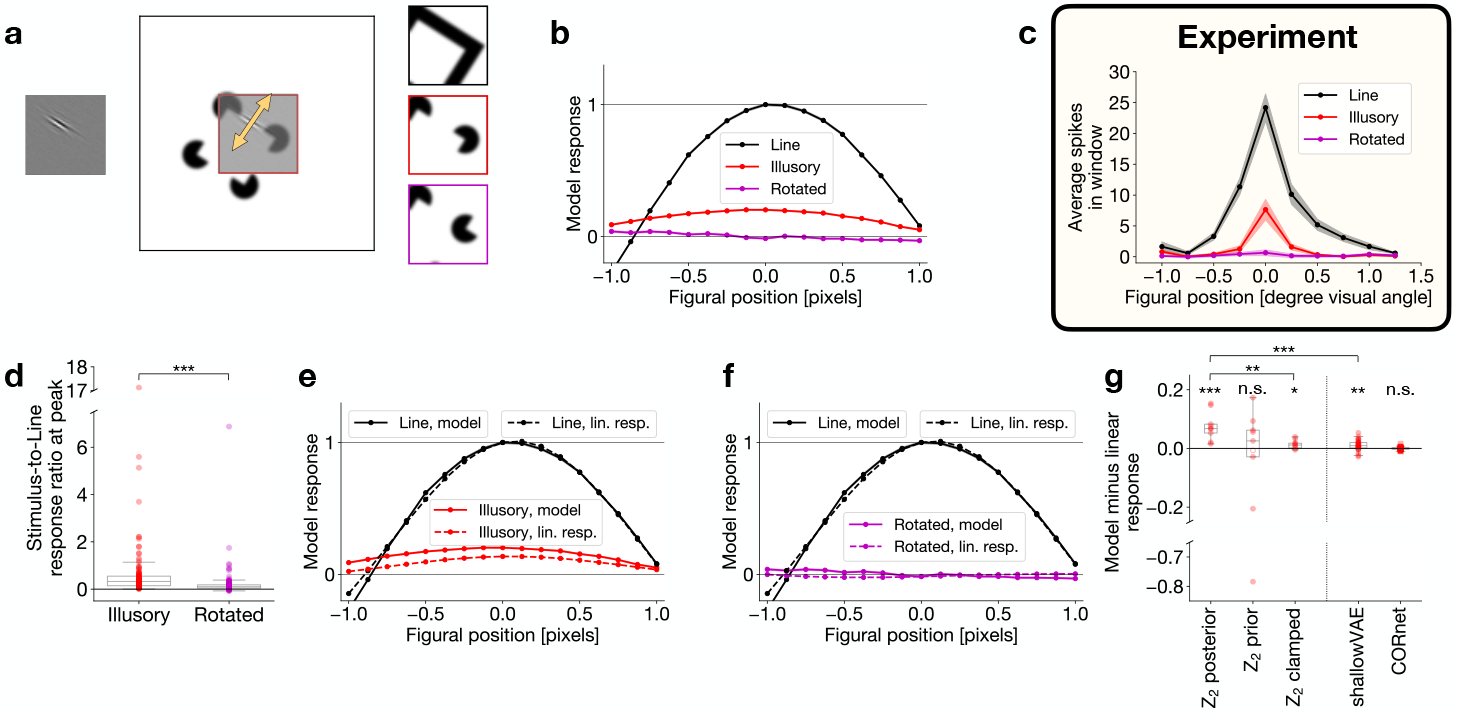
Contribution of top-down computations to illusory contour responses. **a**, Illustration of the illusory contour experiment. *Left*: receptive field of an example model Z_1_ neuron. *Middle*: receptive field-aligned Kanizsa square stimulus. Red border indicates the actual boundary of the stimulus. Arrow: direction of shifts of the stimulus relative to the neural receptive field. *Right*: real square (‘Line’), Kanizsa square (‘Illusory’) and incongruent (‘Rotated’) stimuli. **b**, Mean responses of the Z_1_ neuron in **a** to 500 presentations of the three stimuli as a function of stimulus shift. Shaded regions (invisibly small): s.e.m. **c**, Same as **b** for a selected unit in the V1 of a macaque monkey. Number of trials is unknown. Reproduced from [43] with permission (PNAS, Copyright (2001) National Academy of Sciences, U.S.A.). **d**, Ratio of mean Z_1_ responses to ‘Illusory’ and ‘Line’ stimuli and to ‘Rotated’ and ‘Line’ stimuli, respectively, at the ‘Line’ response peak, for the largest stimulus sizes that fit into the patch, for the analyzed Z_1_ population (*n* = 114, Methods; center line, median; box limits, upper and lower quartiles; whiskers, 1.5× interquartile range). Mean peak ratios: 0.70 (‘Illusory’), 0.23 (‘Rotated’), the latter being significantly less than the former (one-sided paired two-sample ttest, P = 1.6 × 10^−5^, t = 4.3, df = 226). **e**, Same as **b** for the ‘Line’ and ‘Illusory’ stimuli, together with linear responses of the Z_1_ neuron (dashed lines). Magnitude of the linear response to the ‘Line’ stimulus was scaled to the mean response peak to the ‘Line’ stimulus. Shaded regions (invisibly small): s.e.m. **f**, Same as **e** for the ‘Line’ and ‘Rotated’ stimuli. **g**, Differences between the mean and linear responses to the ‘Illusory’ stimulus (center line, median; box limits, upper and lower quartiles; whiskers, 1.5× interquartile range). *Left*: Same restricted Z_1_ population of TDVAE in three different conditions (*n* = 9 each; see text, Methods): intact inference in Z_1_, inference without stimulus-specific information at Z_2_, inference with Z_2_ clamped to zero. *Right*: Illusory responses in two control models: shallow VAE (*n* = 29), and feed-forward goal-directed model (*n* = 40). *Filled circle*: significant boosting or suppression. *Hollow circle*: no significant effect. *Square*: deterministic model.

Using the TDVAE model, we designed a stricter control. Overlap between the receptive field of Z_1_ neurons with the elements of the Kanizsa stimulus could contribute to model responses. To isolate top-down effects, we calculated linear responses of the neurons to the Kanizsa stimuli (Fig. 4e). We found significant boosting of the linear response in TDVAE for the example neuron. By restricting the set of analyzed neurons to those having (1) only a small overlap between the ‘Illusory’ stimulus and the receptive field and (2) all four response peaks in Fig. 4e close to each other (a proxy of regular peak shapes), consistent boosting was observed (*n* = 9; significant boosting: *n* = 9, one-sided one-sample t-test, P < 0.05; population statistics: mean model–linear response difference: 0.076; one-sided one-sample t-test against 0: P = 0.00076, t = 4.7, df = 8) (Fig. 4g, Methods). Incongruent Kanizsa stimuli produced limited and inconsistent modulation in the same set of Z_1_ neurons (*n* = 9; significant boosting: *n* = 5, significant suppression: *n* = 3, one-sided one-sample t-test, P < 0.05; population statistics: mean model– linear response difference: 0.0065; two-sided one-sample t-test against 0: P = 0.39, t = 0.92, df = 8) (Extended Data Fig. 9b). Confirming the contributions of top-down connections to illusory activity, modulation of Z_1_ responses was significantly weaker in a model that lacked Z_2_ than in the TDVAE model (*n* = 29; mean model–linear response difference: 0.011; one-sided one-sample t-test against 0: P = 0.0016, t = 3.2, df = 28; one-sided independent two-sample t-test against TDVAE: P = 2.2 × 10^−7^, t = −6.1, df = 36; Fig. 4g).

To explicitly investigate the contribution of Z_2_ to illusory responses in Z_1_, we manipulated Z_2_ in two different ways. First, to simulate severing feed-forward connections to Z_2_, we removed stimulus information by sampling the prior of Z_2_ instead of the posterior when presenting a Kanizsa stimulus. As predicted, severing feed-forward connections results in TDVAE responses matching linear responses (*n* = 9; mean model–linear response difference: −0.068; two-sided one-sample t-test against 0: P = 0.50, t = −0.71, df = 8; Fig. 4g). Second, as a proxy to severing feedback connections, we simulated illusory contour responses in Z_1_ by clamping Z_2_ responses to zero. This also showed modulations similar to the shallow VAE rather then those of TDVAE (*n* = 9; mean model–linear response difference: 0.014; one-sided one-sample t-test against 0: P = 0.011, t = 2.9, df = 8; one-sided paired two-sample t-test against intact TDVAE: P = 0.0015, t = −4.2, df = 16; Fig. 4g). To show that illusory contour responses are a consequence of learning in the task-general model, we repeated the experiment with the goal-directed feedforward model. No illusory contour responses were observed in this goal-directed model (*n* = 40; mean model–linear response difference: −0.00030; two-sided one-sample t-test against 0: P = 0.71, t = −0.37, df = 39; Fig. 4g).

In non-human primates, extensive evidence supports that stimulus which resonates with the sensitivities of higher-order visual areas produce top-down effects in V1 [44, 45]. A simple analysis of texture-level information in Z_1_ provides a strong indication that statistics represented at Z_2_ flows back to Z_1_ through top-down connections: we performed texture decoding both with intact Z_2_ and with removing stimulus information from it. This analysis confirmed that the small but significant decodability of texture-family information in Z_1_ was a consequence of top-down feedback from Z_2_ (*n* = 5 fits from different random seeds; mean accuracies ± stds: 0.1943 ± 0.0012, 0.0986 ± 0.0025; one-sided independent two-sample t-test: P = 9.6 ×10^−13^, t = 70, df = 8; Fig. 5a). Direct ablation of V2 inputs is currently not widespread in neuroscience research, instead experimental studies use the temporal evolution of V1 signals to distinguish feed-forward and top-down components. We model this effect by assuming that without direct stimulus contribution the cortex samples the prior [12, 33, 46]. Thus, pre-stimulus responses were calculated by sampling the prior of both Z_1_ and Z_2_, while at early-stimulus responses we assumed that stimulus information reaches Z_1_ but Z_2_ responses still rely on the prior. Only late responses showed illusory activity in Z_1_ (*n* = 9 everywhere; two-sided one-sample t-test against 0 everywhere except late Illusory; df = 8 everywhere; pre Illusory: mean: 0.0011, P = 0.49, t = 0.72; pre Rotated: mean: 0.00077, P = 0.65, t = 0.47; early Illusory: mean: 0.068, P = 0.50, t = −0.71; early Rotated: mean: −0.091, P = 0.42, t = −0.85; late Illusory: mean: 0.076, onesided one-sample t-test against 0: P = 0.00076, t = 4.7; late Rotated: mean: 0.0065, P = 0.39, t = 0.92; Fig. 5b), consistent with late-emergence of illusory contour responses in macaques [43]. Similar late-emerging top-down effects were identified when high-level statistics of stimuli was manipulated through phase scrambling (*n* = 5 fits from different random seeds; mean accuracies ± stds: pre: 0.0680 ± 0.0007, early: 0.0986 ± 0.0025, early scrambled: 0.1007 ±0.0012, late: 0.1943 ± 0.0012, late scrambled: 0.1709 ± 0.0020; two-sided independent two-sample t-test between early and early scrambled: P = 0.15, t = −1.6, df = 8; one-sided independent twosample t-test between late and late scrambled: P = 1.8 × 10^−8^, t = 20.3, df = 8; Fig. 5c) and when large-scale integration of visual elements was disrupted by manipulating contours (*n* = 57 in both cases; early: mean effect size: −0.080, two-sided one-sample t-test against 0: P = 0.52, t = −0.65, df = 56; late: mean effect size: 0.36, one-sided one-sample t-test against 0: P = 0.00014, t = 3.9, df = 56; Fig. 5d, Extended Data Figs. 9c,d). These results are confirming results obtained from electrophysiology recordings from V1 of macaques when presenting stimuli matching those used in our experiments [44, 45].

**Fig. 5.**
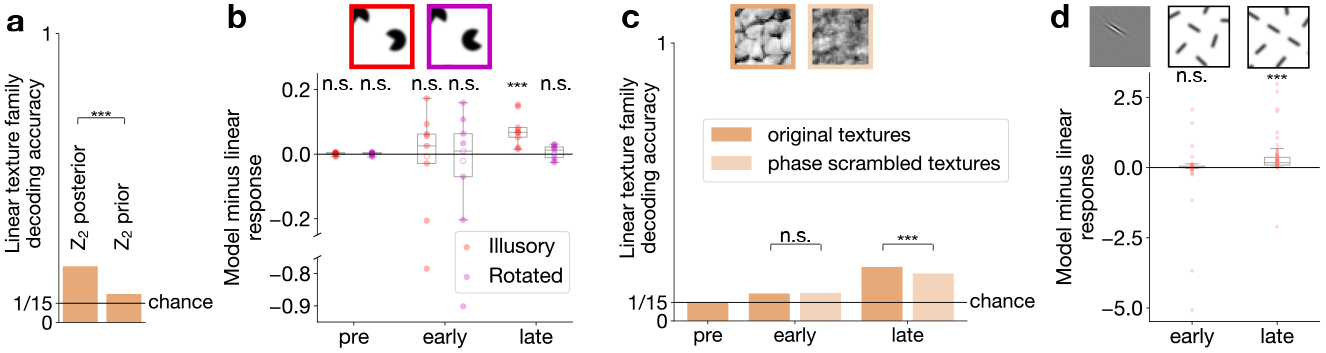
Top-down influences shaping the representation in Z_1_. **a**, Performance of texture family decoder from mean responses of Z_1_ with intact inference, and without stimulus-specific information at Z_2_. Means of *n* = 5 fits are shown; s.d. < 0.01 everywhere. **b**, Qualitative time course of illusory contour responses in the Z_1_ population from Fig. 4g (*n* = 9; center line, median; box limits, upper and lower quartiles; whiskers, 1.5× interquartile range). *Inset*: example stimuli. **c**, Qualitative time course of texture family decoding performance from intact and phase-scrambled versions of textures. Means of *n* = 5 fits are shown; s.d. < 0.01 everywhere. *Inset*: example stimuli. **d**, Qualitative time course of contour completion experiment. *Top*: Illustration of stimuli. Optimally oriented, identical line segment is shown in the receptive field of a Z_1_ neuron (*left*) under the condition that the neighboring visual field is filled with randomly oriented segments (*center*) or the neighboring segments continue the segment that covers the receptive field (*right*). *Bottom*: Qualitative time course of the response intensity difference between the contour completion and random conditions in the Z_1_ population (*n* = 57; center line, median; box limits, upper and lower quartiles; whiskers, 1.5× interquartile range).

**Fig. 6.**
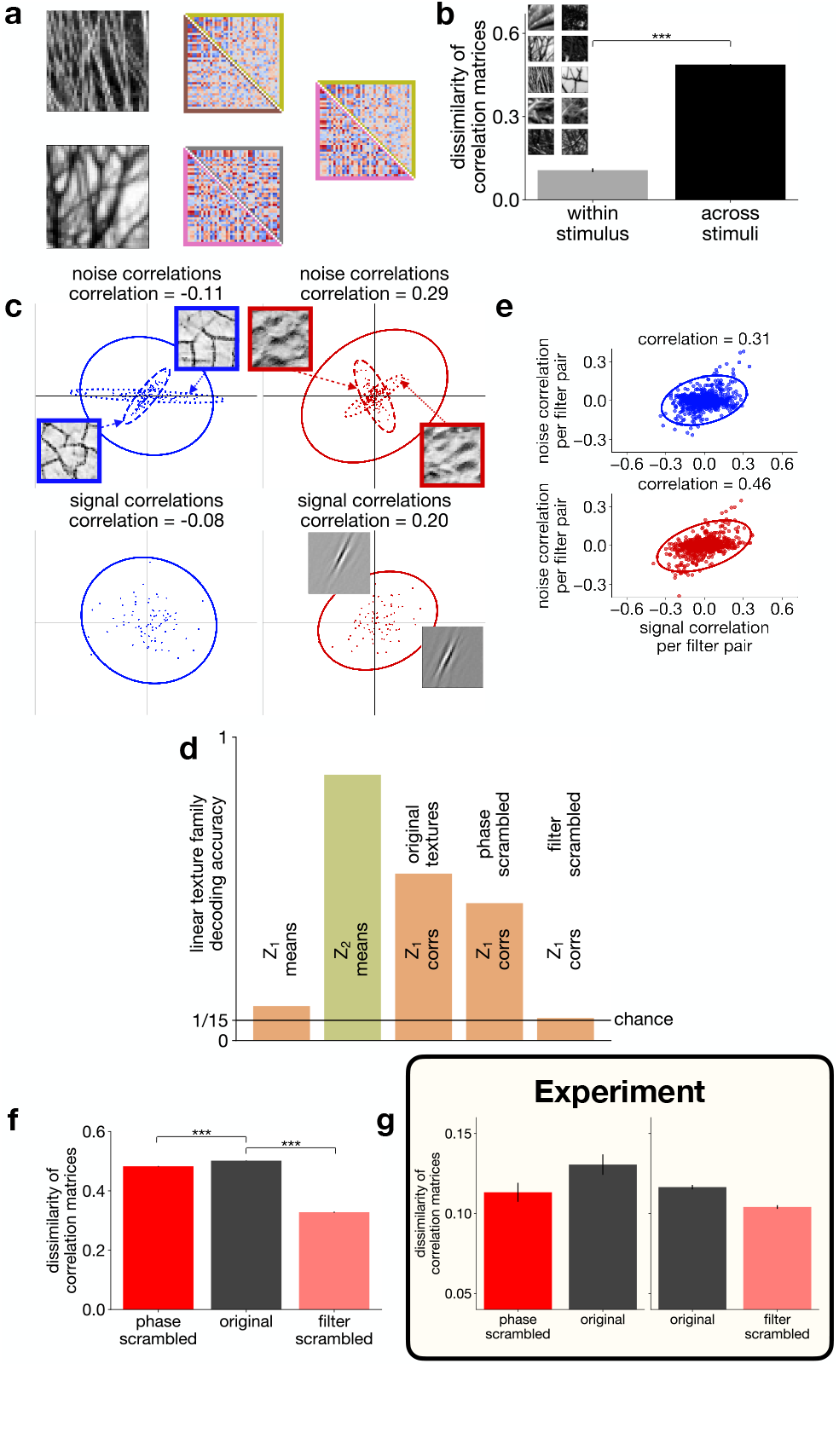
Contribution of top-down influences to response correlations in Z_1_. **a**, Z_1_ noise correlations for *n* = 40 neurons with central, localized, medium wavelength receptive fields for two example natural images (*left*). *Center*: Effect of sampling noise. Comparison of noise correlations from two independent sets of trials (*n* = 80 each, *lower* and *upper triangles*, respectively). *Right*: across-image noise correlation differences. Data taken from the central column (*border colors* matching those in the middle column). **b**, Within- and across-image dissimilarity of noise correlations (from *n* = 80 trials each), averaged over 10 images (*inset*). **c**, *Top*: Response covariance of two Z_1_ neurons to individual images. *Dashed* and *dotted covariance ellipses* correspond to the two *inset images. Solid ellipses*: average covariance ellipses for *n* = 80 images per texture family. *Left* and *right columns*: two example texture families. *Text labels*: per-family mean noise correlation coefficients. *Bottom*: Trial-averaged mean responses of the same Z_1_ units to *n* = 80 images (dots) from the same texture family as top. *Covariance ellipses, text labels*: signal correlations calculated from these responses. **d**, Texture family decoding performance from Z_1_ (*n* = 100) and Z_2_ (*n* = 6) mean responses (as in Fig. 2d) and Z_1_ noise correlations (from the same *n* = 100 Z_1_ units as for Z_1_ means). Decoders were constructed for texture images, as well as phase and filter scrambled versions of these. Bar heights: means for *n* = 5 random seeds; s.d. *<* 0.01 everywhere. *Black line*: chance performance. **e**, Relationship between noise and signal correlations for a population of Z_1_ (*n* = 40 Z_1_ units) neuron pairs (*dots*) for two texture families. Colors and data as in panel **c. f**, Across-stimulus dissimilarity of noise correlations (20 natural images, four noise correlation matrices per image, each calculated from 80 trials). Error bars: s.e.m. **g**, Same as **f** in V1 multiunit recordings from macaque monkeys. The two comparisons are calculated from data recorded in distinct experimental sessions. Reproduced from [47] with permission (PNAS).

### Response correlations emerging through contextual priors

Illusory contour, contour integration, and phase scrambling experiments demonstrate how contextual priors affect V1 neurons through features represented higher in the cortical hierarchy, V2 in our specific case. These contextual priors carry information about the regularities of the surroundings of a neuron’s receptive field, such as the tendency of another neuron to be jointly active in the local environment. These regularities in the local environment can change from image to image and thus the changing contextual prior dictates different co-activation patterns between neurons (Figs. 6a,b, Extended Data Fig. 10a). Such stimulus-specific co-activation patterns are captured in correlations between neuronal responses to individual images and thus noise correlations. Indeed, across-stimulus dissimilarity of noise correlations is larger than that expected from estimating noise correlations from a finite sample (mean dissimilarities: 0.487 and 0.107, respectively; one-sided independent two-sample t-test, P = 3.4 × 10^−91^, t = −38.4, df = 188). Noise correlations provide us with a tool to identify and characterize top-down influences as contextual priors contribute to these. Importantly, if regularities in a set of images are similar, the contextual prior and thus the noise correlations are not changing, therefore texture family-specific activity in V2 is expected to induce texture family-specific noise correlations in V1 (Fig. 6c).

We measured the texture-family specificity of noise correlations in the TDVAE model by constructing linear decoders that we applied to noise correlation matrices of a small Z_1_ neuron population (Fig. 6d). In the TDVAE, it is the top-down influence that is exclusively responsible for the covariability of Z_1_ neurons as the variational posterior is correlation-free. To disentangle top-down correlations from private variability, we use the changes in the mean of the posterior of Z_1_, which is resulting from sampling Z_2_, to assess correlations. Additional private variability reduces the magnitude of noise correlations but it does not affect the overall trends reported in this section. As Z_2_ response means show linear decodability of texture family information, linear decoding of texture family from Z_1_ correlations was also highly effective (Fig. 6d). In addition, stimulus manipulations weakening the texture representation in Z_2_ like phase and filter scrambling also reduce the texture family decodability from Z_1_ correlations (Fig. 6d). In contrast to Z_1_ response correlations, Z_1_ response means are worse suited to linear decoding of texture family information (Fig. 6d).

The contextual prior reflects the regularities of the environment that are specific to the particular context. For texture families, the tendency of a pair of neurons to be jointly active for different texture images, referred to as signal correlation, is expected to be captured by the contextual prior of a well trained model. Indeed, Karklin and Lewicki argued that the correlations between mean filter responses is characteristic to texture families [48], and we also found that such texture family-specific signal correlations were present between an example pair of neurons (Fig. 6c). Thus, the contextual prior is related to two different forms of joint response statistics: the signal correlations, measuring correlations between mean responses across images, and the noise correlations, measuring correlations between neurons following multiple presentations of the same image. Therefore, we measured for every texture family the relationship between these two quantities for a large number of Z_1_ neuron pairs. We found strong dependence of noise correlations on signal correlations in the responses of pairs of Z_1_ neurons (Fig. 6e, Extended Data Fig. 10b).

We test the consequences of the proposal that noise correlations are shaped by contextual priors in a modified setting. We argued that stimulus-specific noise correlations arise due to top-down feedback that carries information about high-level inferences. Consequently, removal of high-level structure is expected to result in reduced stimulus-specificity of noise correlations in Z_1_. Both our earlier analyses (Figs. 4, 5) and electrophysiology data [44] support the idea that texture-statistics related information is delivered to Z_1_ through top-down feedback from Z_2_. The former analysis concerned top-down induced changes in response mean in Z_1_ neurons and here we extend this to top-down influences on noise correlations. To test this, we used the same stimulus manipulations that we used for exploring the properties of Z_1_ and Z_2_ representations: phase and filter scrambling of natural stimuli (Methods). We calculated the dissimilarity of noise correlation matrices for natural images and for phase- and filter-scrambled versions of the same natural images (Methods). We found reduced dissimilarity for phase scrambled versions of natural images (Fig. 6f, mean for intact images: 0.502, mean for phase scrambled images: 0.483, one-sided paired two-sample t-test, P = 9.0 × 10^−82^, t = 19.7, df = 6078) and filter scrambled versions (mean for intact images: 0.502, mean for filter scrambled images: 0.328, one-sided paired two-sample t-test, P *<* 10^−100^, t = 92.7, df = 6078). Similar reduction in the dissimilarity of noise correlations for phase- and filter-scrambled versions of natural images was found in the V1 of macaques [47] (Fig. 6g). Note, that larger magnitude of dissimilarity, i.e. higher stimulus specificity of noise correlation, in the model is a direct consequence of focusing on quantifying correlated variability and removing private variability from analyses. Sampling the posterior of Z_1_ reduces the measured stimulus specificity of noise correlations (Extended Data Fig. 10c,d), making the magnitude comparable to the experimentally obtained values. The exact level of reduction depends on the details of how sampling is performed, as well as experiment details, including trial length. Given the 400-ms time window of the experiments in [47], we have explored a number of variants in sampling (Extended Data Fig. 10c,d). While systematic changes were introduced by the relative number of samples from Z_1_ and Z_2_, relative differences of correlation dissimilarities across stimulus types were consistent. In summary, similar stimulus statistics dependence of noise correlations in Z_1_ of TDVAE and V1 of macaques provide support that contextual priors emerge as a consequence of hierarchical inference.

## Discussion

In this paper we extended deep learning models of the ventral stream by developing a deep generative model. The developed TDVAE model departs from the feed-forward architectures of earlier deep learning accounts as it naturally accommodates top-down interactions: top-down connections serve the implementation of contextual priors for hierarchical inference. Importantly, to investigate how neural representations emerge and shape computations, our normative model was end-to-end trained on natural images alone and neural representations could be studied in a setting where no free parameters were available. TDVAE displayed strong alignment with the texture representations identified in recordings from V1 and V2 of non-human primates and rodents. Our deep generative model was shown to reproduce key experimental signatures of top-down computations. Importantly, studying the computations in TDVAE enabled us to gauge how feed-forward and top-down contributions interact in V1. Our results highlight that sensitivities of neurons in V1 are shaped by top-down influences and the same principles also explain how top-down influences contribute to stimulus-dependence of noise correlations in V1. Beyond reproducing a wide range of experimental findings, we demonstrate that the framework opens a new window on exploring the representation, computations, and the circuitry of the early visual system.

Generative models have long been implicated in perceptual processes as these establish an essential framework for learning in an unsupervised manner. Conceptually, generative models summarize our knowledge about the world and describe the way observations are produced as a result of the interactions between (a hierarchical set of) features. Perception is assumed to invert the generative process by inferring the features that underlie observations [9, 32]. However, modeling tools to investigate neural processes have been limited. Significant contributions to understanding the computations in the visual cortex were made with non-hierarchical, linear models [29], manually designed hierarchical extensions [49, 50] and linear hierarchical models [51, 52]. Non-hierarchical extension of the linear models were shown to account for extra-classical receptive field effects and response variability [36, 37, 53, 49]. Further, a mild hierarchical extension that explicitly modeled dependencies between features represented by receptive fields in natural images [54], when applied to V1, could account for complex cell properties [48]. Here we propose to overcome the obstacle of training models that are both hierarchical and flexible enough to accommodate the complexity of natural images by recruiting the recently developed Variational Autoencoders (VAEs) from machine learning [25, 26] and extending these towards hierarchical computations [27, 28]. Non-hierarchical versions of VAE have previously been applied to various stages of the ventral stream, including early visual cortex [55, 30] and inferotempral cortex [56]. Interestingly, a hierarchical VAE proved useful for modeling computations along the dorsal stream [57] and could establish relationships to responses in MT but top-down computations were not addressed.

The proposed TDVAE architecture relies on a limited set of assumptions while remaining as faithful as possible to the principles of probabilistic inference. In order to stick with the guarantees of VAEs, we did not rely on extensions of the framework that encourage disentanglement [56, 58], regularization methods that might distort the learned representations (see Methods), or methods that help limiting the number of learned parameters (convolution). TDVAE implements an approximate form of probabilistic inference, which we related to neural response statistics through the sampling hypothesis. A range of studies delivered support that stochastic sampling accounts for patterns in response variability in V1 and beyond [59, 36, 60, 7, 61, 62, 12, 63, 37]. In our study we obtained samples from the variational posterior of the recognition model and performed this sampling with a fixed neural architecture, yielding the so-called amortized inference. Alternatively, a sampler could be constructed that obtained samples directly from the generative model. We motivate the chosen approach by three arguments. First, the variational posterior provided a useful tool to disentangle the contribution of top-down influences to response correlations. Second, by keeping our approach as close as possible to standard machine learning approaches, the framework can be naturally extended in further directions. Third, we found the circuitry implied by the recognition model to be a good inspiration for interpreting layer-by-layer computations. In our study, an important corollary of sampling was that it enabled the model to represent correlated posteriors. These correlated posteriors resulted directly from top-down contextual effect. Still, the complexity of the true posterior was only approximated through this contextual effect and a more flexible posterior could be obtained from direct samples from the generative model. Such more flexible sampling of the posterior has the potential to provide insights into other forms of the response statistics of neuron populations recorded with high-throughput techniques. How neural circuits implement these computations is a critical question. Stabilized supralinear networks were shown to produce response statistics with well controlled correlation and variance patterns [64]. Combining amortized inference and sampling to perform inference can provide insights into the recurrent interactions within the lateral structure of V1 [65, 63] but a version that could perform hierarchical computations has not been explored so far.

The deep generative modeling approach presented here complements the more traditional works on generative models of vision that directly rely on statistical insights to construct the neuronal circuitry [53, 66]. While the multi-layer perceptron networks that implement the computational components of the recognition and generative models in TDVAE are endowed with the flexibility necessary for high-complexity models, these prevent obtaining direct mechanistic insights that more traditional approaches offer [66]. These mechanistic insights might help to establish inductive biases for deep generative models for more efficient learning of natural image statistics [67]. Note that the specific form of the recognition model has a consequence that feedback to V1 in our model is implemented through a single-step top-down pass, avoiding highly recurrent refinement of the responses between V1 and V2. Experiments relying on time delays between neural responses in V1 and V2 are compatible with our account but more precise experimental data can identify more elaborate recurrent computations.

Top-down interactions have previously been implicated in an alternative computational framework to our deep generative models, predictive coding [68, 69]. The two frameworks have strong parallels: both are unsupervised, rely on a generative model, and are hierarchical. The two frameworks can be distinguished based on the goal top-down connections serve. While in our account, top-down interactions contribute to rich inferences that can serve flexible task execution [23, 10], in the predictive coding framework top-down connections serve an efficient representation of the stimuli [70]. In our framework inference is inherently probabilistic. Probabilistic inference augments classical inference frameworks by representing the uncertainty associated with the interpretation of the incoming stimuli [12, 71]. Theoretical arguments provide strong support for probabilistic inference and have also gained strong empirical support from behavioral studies [11, 72]. More importantly, the representation of uncertainty has been identified in neuronal responses in V1 [71] and such probabilistic computations have explained a wide range of phenomena [36, 37, 59, 63]. Representation of uncertainty is not part of the existing formulations of predictive coding models (inference is performed at the maximum a posteriori level [73]), albeit proposals have been made to extend the original framework of predictive coding in this direction [74]. In practice, our results on stimulus-dependent correlations are tightly linked to a probabilistic representation and are therefore beyond the scope of predictive coding accounts.

In the case of TDVAE, the computational graph of the recognition model provides insights into the potential circuitry serving the computations. Feed-forward information reached Z_2_ through a relay from Z_1_, which can be implemented by layer 4 neurons. Feedback from Z_2_ can be interpreted as projections to the superficial layers of V1, which is supported both by anatomical data [75] and the progression of texture information across laminae [44]. Recent experiments have highlighted that so-called feedback receptive fields in layer 2/3 are inherited through top-down interactions, and these are specific to layer 2/3 but are absent from layer 4 [76, 77]. Accordingly, we hypothesize that layer 2/3 of V1 could accommodate Z_1_, which integrates feed-forward and feedback information. Indeed, we found that texture sensitivity in Z_1_ was inherited from higher processing layers. Note, that the neural networks that implemented the components of TDVAE (Z_1_-ff, Z_2_-ff, Z_1_-INT, Z_1_-TD) were considered as general computation elements. Therefore we made no attempts to link constituent neurons to the neurons of the ventral stream but this can be a potential subject of future research.

Top-down computations represent one specific form of recurrence in the visual cortical computations. Within-area inhibitory and excitatory recurrence is characteristic of the cortex. Such recurrent interactions have been investigated through generative models of natural images in the context of modeling collective modulations of neuronal responses by stimulus contrast, which were shown to contribute to nonlinear response properties of V1 neurons [53, 37, 49]. Interestingly, this neurally well-motivated generative model has recently been shown to extend and improve the variational autoencoder framework [67], which can also be integrated in the future with the hierarchical VAE framework discussed here. Another role of recurrence has been proposed for performing fast and accurate inference in generative models of natural images by neural networks [63]. Recurrent computations have gained traction in deep discriminative models too and were shown to surpass purely feed-forward models in prediction of neural activity [78]. More specifically, recurrence was shown to contribute to such predictions when stimuli were perceptually challenging [5]. This result is in line with our argument that recurrent activity helps under perceptual uncertainty. Indeed, in human MEG recordings, contribution of recurrent computations was shown to be important when processing occlusion, another source of perceptual uncertainty [79]. This provides an insight that beyond the uncertainty arising through the hierarchical structure of natural images, uncertainty arising from the interactions of different visual elements might rely on recurrent instead of top-down computations. Generative models that combine hierarchical and as well as lateral interactions have been proposed in machine learning [80], which can provide inspirations for further extensions We emphasized the role of top-down interactions in inferring perceptual variables through establishing contextual priors. Importantly, priors that are not dependent on a context can be efficiently implemented without the need of top-down connections [81, 82, 83]. The advantage of top-down induced priors is their flexibility as these can depend on the wider context and such can depend on the stimulus itself. In contrast with the implication of top-down connections in perceptual inference, a wide array of studies consider attentional modulation as an important factor contributing to top-down interactions [84]. Reconciling these alternative interpretations is a critical challenge. We argue that considering task-relevant variables besides perceptual variables can be interpreted in the same inference framework [62]. Indeed, top-down delivered information in the early visual cortex was shown to be specific to the task being performed in humans [85]. Also, it has been argued that attentional effects can rely on a similar computational scheme, in particular hierarchical inference, to that proposed here for perceptual inference [60, 86]. Interestingly, the top-down modulation of noise correlation patterns has also been demonstrated under changing task conditions [87]. The question how perceptual and task-related variables are integrated in the circuitry that establishes top-down interactions remains a largely open question but recent studies indicate that they may share the same communication pathways [88].

We argued that the emergence of illusions can distinguish goal-directed models with essentially feed-forward architectures from task-independent models that perform probabilistic inference through establishing contextual priors. Indeed, a variety of perceptual illusions have been reported to be at odds with goal-directed models [89] and these have also been challenged by higher-level perceptual effects [90]. Interestingly, modeling works, which had a focus on feedback but not on the cortical representation, highlighted an additional perceptual effect related to the Kanizsa illusion that relied on higher-level shape information [91, 92]. This is in line with reports that cortical areas beyond V2 are central to the Kanizsa effect [93]. Indeed, higher-level features that are represented beyond V2 could contribute to top-down boosting of illusory responses, but the critical contribution of texture-representing region became evident through the evidence that causal intervention in the lateromedial region of mice eliminated illusory responses [94]. Hierarchical Bayesian models have been proposed to account for illusory contour responses [95, 96], which implicate recurrent connections from higher visual areas. While these hand-crafted models can capture the signatures of the Kanizsa experiment, it is an end-to-end trained model that can account for the interplay between the learned representations of a neural network and the computations taking place in them. While illusions are key signatures of the contextual priors that emerge through top-down interactions, these interactions have wider ranging effects. Recently, modulation of V1 neurons by the visual context of their receptive field has received support from a hierarchical inference account [50]. Disentangling the contributions of feed-forward and feedback components is thus a key challenge of systems neuroscience but novel tools promise to break this impasse by providing a quantitative testbed for theories [97].

In our study we found a robust texture representation in a layer homologous to V2. In our approach we did not encourage learning texture-like representations since the model was trained on natural images. We argue that multiple factors contributes to the properties of the representations emerging at different layers of the hierarchy. In the TDVAE, low-level representation was constrained by the inductive bias applied to the generative model. Interestingly, this inductive bias also implicitly constrained the recognition model: contraintutively, we found that texture decoding performance from Z_1_ was higher in an untrained model than from the natural image trained model, indicating that the nonlinearity required to capture texture-like statistics is ‘unlearned’. Texture statistics are well characterized by a spectrum of second-order statistics [40]. Therefore a limited computational complexity model can capture textures. Thus, we expect that a generative model with a deeper hierarchy but with a generative layer that is still constrained to second-order statistics will learn similar texture representation. Higher levels of the hierarchy that can recruit further nonlinearities will be able to capture more nonlinear features. We found similar representations in variants that featured a more complex generative model. We argue that this might highlight the contribution of another factor: image size. The statistics relevant to textures have a characteristic scale. Intuitively, more complex features might have statistical signatures spanning even larger spatial scales. We speculate that the limited patch size actually affected the features TDVAE developed sensitivity to. Taken together, our results demonstrate a very robust texture representation that emerges for natural images in a nonlinear generative model. The constraints implicit in our implementation of TDVAE contributed to this robustness, and provide the necessary insights to build generative models with deeper hierarchies that capture more complex features and higher-level processing in the visual cortex. Several machine learning studies indicate that developing multiple levels of hierarchy is a viable route with VAEs [27, 28]. The two-layer TDVAE can be extended to higher levels of the hierarchy. Distinguishing the computations at different layers of the hierarchy can be achieved by inductive biases (similar to the linear limitation we implemented in one component of the generative model) or explicit architectural constraints (such as limiting the statistics by limiting the visual field a neuron has access to [98]). The nature of the inductive biases shaping higher level representations is subject to further research.

## Methods

### Generative models

We study Markovian two-layer hierarchical latent variable generative models that learn the joint distribution of observations and latent variables in the form

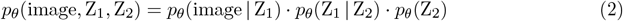

where image denotes the pixel intensities of the modelled images, Z_1_ and Z_2_ denote stochastic model layers modeling biological neurons in visual cortical areas V1 and V2, respectively, and *θ* denotes the parameters of the generative model. To make inference tractable, we train Variational Autoencoders (VAEs) [25, 99] which supplement the above generative model with a separate recognition model *q*_Φ_(Z_1_, Z_2_ | image) with parameters Φ that establishes a variational approximation of the true posterior distribution *p*_*θ*_(Z_1_, Z_2_ | image).

In general, the tractable Expectation Lower Bound (ELBO) for a two-layer VAE can be written as

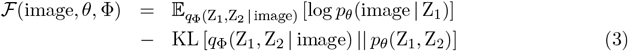

where KL [·||·] denotes the Kullback–Leibler divergence of two probability distributions.

In our main model, TDVAE, we factorize the variational posterior in the following way:

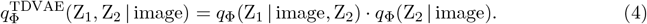

This leads to the recognition model architecture depicted in Fig. 1f and the following objective function (ELBO):

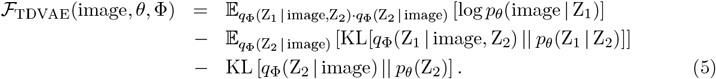

In our alternative model ffVAE the top-down interaction is not manifested in the factorization of the variational posterior:

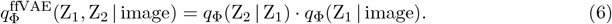

This results in the following objective function:

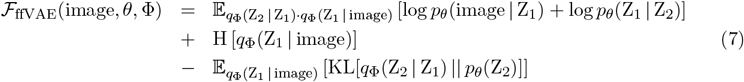

where H [·] denotes the entropy of a probability distribution.

For baseline we also trained a non-hierarchical VAE which we call shallow-VAE and has the objective function

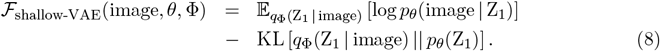

For clarity, in the following we will omit *θ* and Φ from the notation of the generative and recognition models, respectively.

### Architectural details

Inspired by models of V1 activity [29], we chose to implement the Z_1_ layer of latent variables as an overcomplete layer (dim(Z_1_) = 1800 *>* dim(image) = 40^2^ = 1600) with a sparse (Laplace distribution) prior that is in a linear generative relationship to observations: 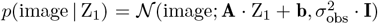 where *σ*_obs_ denotes the standard deviation of the independent Gaussian observation noise for which we used *σ*_obs_ = 0.4 in each presented model. While the dimensionality of Z_1_ permitted learning an overcomplete representation, in practice the dimension of the learned representation (1256) was complete, such that the number of actively participating model neurons was equal to the number of independent dimensions (after the whitening of the images). In addition, we chose dim(Z_2_) = 250 for the hierarchical models TDVAE and ffVAE. While this may seem like an artificially induced compression constraint compared to the 1800 units in Z_1_, you will see in Table 4 that 250 dimensions were enough to accommodate the entirety of the learned representations in Z_2_.

The components of the generative and recognition models were parameterized through neural networks with the softplus activation function. We used fully connected networks (multilayer perceptrons, MLPs) instead of convolutional networks (CNNs) to avoid the indirect effects of CNNs on emerging representations. Extended Data Table 2 highlights the number of hidden units for each part of the computational graph in the shallow-VAE and the ffVAE models.

The computational graph of the recognition model for the TDVAE model (depicted in Extended Data Fig. 1) is nontrivial and contains further inductive biases. There are four MLPs defined for the recognition model as depicted in Fig. 1f. The first neural network Z_1_-ff maps the pixel space to a layer L_x_ that is shared between the computations of *q*(Z_2_ | image) and *q*(Z_1_ | image, Z_2_). From L_x_, a second MLP (Z_2_-ff) computes the means and variances of the *q*(Z_2_ | image) distribution. The third MLP, Z_1_-TD transforms Z_2_ into a layer L_z_. We fuse the information from image and Z_2_ by concatenating L_x_ and L_z_ and applying an MLP Z_1_-INT to the concatenated layer to obtain the means and variances of *q*(Z_1_ | image, Z_2_). The number of hidden layers and hidden units used in each MLP to calculate the means and standard deviations of the conditional generative and variational posterior distributions is shown in Extended Data Table 3 for the different TDVAE models referenced in the paper.

As detailed in Figs. 4, 5 and the accompanying text, 40×40-pixel image patches proved to be too small for studying certain top-down effects with the TDVAE model family. For this reason, we created two larger, 50×50-pixel models called shallow-VAE-50px and TDVAE-50px that are proportionally upscaled versions of the 40×40-pixel shallow-VAE and TDVAE models, respectively (see Extended Data Tables 2, 3, 4, and 5). The qualitative properties of the upscaled TDVAE models match the 40×40-pixel models, so we omit them from the discussion below. In the paper, we use the 50×50-pixel models only for the illusory contour and contour completion experiments; all other results were reached with the 40×40-pixel models.

Finally, we assessed the potential effect of the shared Z_1_-ff mapping by comparing a standard 20×20-pixel TDVAE model with shared encoding called TDVAE-20px and a corresponding ‘TDVAE-20px non-shared’ model (for model parameters, see Exteded Data Table 3). In the latter, Z_2_-ff took its input from the input image instead of the output of Z_1_-ff, and its MLP layers were prepended with an MLP with the same dimensions as Z_1_-ff (for these notations, see Fig. 1f).

### Relating inference in the generative model to neural activity

Features learned by the model are identified with the response properties of individual latent variables, termed model neurons, of the recognition model (Eq. 4). Intensity of response strength of the model is identified with the intensity of the neuronal responses. As the generative modeling framework yields a probabilistic representation, we need to adopt a representational scheme to represent these probabilities. We adopt the sampling hypothesis [59, 33, 10], in which neural activity in any given moment in time represents a sample from this probability distribution and the sequence of samples yields a histogram of potential values from the distribution. The histogram is an approximate representation of the probability distribution, which can converge to the exact distribution as more samples are collected. This means that neural activity yields progressively better representation of the distribution as time passes. In biological neurons, an approximation of the time in which new samples are obtained can be estimated through the time constant of the response autocorrelation, which is approximately 20 ms [34] in the V1. Thus, in a trial of 500 ms 25 samples can be obtained, and a firing rate in this time window informs the experimenter about the mean of the samples. Trial averages of firing rates provide a more accurate estimate of the mean of the posterior distribution, while response variance carries information about the width of the posterior, i.e. the uncertainty of the estimate.

In contrast with firing rates, Z_1_ and Z_2_ can take positive as well as negative values. Optionally, the values of Z_1_ and Z_2_ can be identified with membrane potentials and thus the recorded firing rates can be considered as membrane potentials fed through a firing rate nonlinearity. In line with earlier works with VAEs [56], we chose to report the values of the latent variables instead of such firing-rate transformed versions in order to increase the transparency of the modeling approach. Earlier works have demonstrated that both membrane potential and firing rate predictions of a generative model of natural images are in line with neural data on systematic changes in response mean and response variance [36], and therefore we focus on non-transformed values of model neuron activities. The firing rate transformation introduces a nonlinearity in responses, which can have substantial consequences on reading the information encoded by neurons [100, 38, 47]. While these affect the results quantitatively, the qualitative trends that our paper is focussing on remain unaffected by such change.

Noise correlations in neural responses are identified in the model with correlations in *q*(Z_1_ | image).

In the specific VAE framework the variational posterior of Z_1_, *q*(Z_1_ | image, Z_2_) is not correlated, thus the source of correlation is marginalization over Z_2_: variance in *q*(Z_2_ | image) results in correlated changes in Z_1_. In practice, instead of fully integrating Z_2_, we sample its variational posterior *q*(Z_2_ | image) and perform Monte Carlo integral:

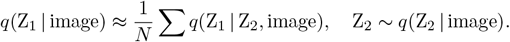

Limiting the source of correlations to result from variance in Z_2_ might seem as a limitation of the model, as the true posterior of Z_1_ might feature other statistical structure too. We argue that this setting actually provides clarity in interpreting how noise correlations emerge in the model and thus how top-down effects shape noise correlations.

### Model training details

We trained all our models with the Adam optimizer on natural image patches with a minibatch size of 128. We found that while the learned Z_1_ representation was robust against the tested regularization techniques (weight decay, gradient clipping[101], gradient skipping[27]), the learned Z_2_ representation was sensitive to them. To eliminate such regularization artifacts, we turned off weight decay and increased gradient clipping and skipping thresholds to have an activation frequency below 10^−6^.

For training our hierarchical VAE models TDVAE and ffVAE we applied *β* annealing, which is a standard technique for training VAEs. This means that in their ELBO losses (5) and (7) we introduced a scaling factor *β* in front of the last term. We started training with *β* = 0.1 for 1000 epochs and then linearly increased *β* to 1 over the course of 1800 epochs and then continued training with the original ELBO loss. No *β* annealing was used for the shallow-VAE model.

We found that our two-layer models TDVAE and ffVAE achieved significantly better ELBO values if we changed the learning rate parameter *α* whenever the validation loss reached a minimum. The following learning rate schedules were found to lead to the highest ELBO values (these are the models reported throughout this paper):

**shallow-VAE** *α* = 3·10^−5^ for 12,000 epochs

**TDVAE linear integration** *α* = 10^−5^ for 40,500 epochs, then 2·10^−5^ for 2100 epochs, and

2.85·10^−5^ for 630 epochs

**ffVAE shallow nonlinearity** *α* = 10^−5^ for 7400 epochs, then 2*·* 10^−5^ for 600 epochs, 3 *·*10^−5^ for 320 epochs, and 10^−6^ for 10 epochs

**TDVAE shallow nonlinearity** *α* = 5·10^−5^ for 4500 epochs, then 2.5·10^−5^ for 9000 epochs

**TDVAE deep nonlinearity** *α* = 10^−5^ for 205,700 epochs, then 2·10^−5^ for 2100 epochs, 3·10^−5^ for 5700 epochs, and 5·10^−5^ for 900 epochs

**shallow-VAE-50px** *α* = 1.5·10^−5^ for 72,000 epochs

**TDVAE-50px shallow nonlinearity** *α* = 10^−5^ for 100,500 epochs, then 3·10^−5^ for 800 epochs. In this case, changing the learning rate alone could not compress the Z_2_ representation to include only texture family encoding units. We promoted compression by increasing *β* to 8.5 and then lowering it back to 1.

**TDVAE-20px** *α* = 10^−4^ for 6000 epochs, then 2 *·*10^−4^ for 1050 epochs, and 3*·* 10^−4^ for 300 epochs

**TDVAE-20px non-shared** *α* = 10^−4^ for 6000 epochs, then 2·10^−4^ for 1050 epochs, 3·10^−4^ for 750 epochs, and 3.4·10^−4^ for 450 epochs In the paper, we present one instance of each model type. The models were trained on an Nvidia GeForce RTX 3080 Ti GPU.

Validation ELBOs at the end of training and the number of active units in the learned representations in Z_1_ and Z_2_ are presented in Extended Data Table 4. Of all presented 40×40-pixel models, TDVAE with a deep nonlinearity in the higher level of its generative model *p*_*θ*_(Z_1_ | Z_2_) reached the highest ELBO value, therefore it is the primary interest of this paper. For all models, Z_1_ learned a complete linear basis of the training data (see below on the effective dimensionality of the natural training data) while the nonlinear representations learned by Z_2_ had different dimensionalities for different models.

### Discriminative model benchmark

When comparing our VAEs to a discriminative model we picked CORnet-Z [41] as a benchmark. This is a purely feedforward model which was optimized for object recognition. It has various blocks which were shown to have high representational similarity to V1, V2, V4, and IT. We used only the ‘V1’ and ‘V2’ blocks. A block in the model consists of two layers, a ‘conv’ and an ‘output’ layer, where the latter is calculated with pooling operations from the former. We found with reverse correlation experiments that the ‘conv’ layer has features reminiscent of Gábor filters (similar to our models’ lower layer) while the output layer has more complex receptive field structure. Therefore we used the ‘conv’ layers to compare CORnet feature values to our model. The model expects 224×224-pixel RGB images as input. In our analysis we only fed non-whitened images to the model where only the blue channel had non-trivial pixel values.

### Natural image datasets

All presented models were trained on a ‘training’ natural image dataset compiled by the authors in a way similar to [30]. We took 320,000 random 40×40-pixel crops, 160,000 random 50×50-pixel crops, or 640,000 random 20×20-pixel crops from the first 1000 images of the van Hateren natural image dataset [102], matched their grand total pixel intensity histogram to the standard normal distribution, and applied ZCA whitening while keeping only 25*π*% of the largest-eigenvalue PCA components. This made the effective dataset dimension 1256, 1963, or 314 instead of the number of pixels in an image patch, 1600, 2500, or 400, respectively. This whitening procedure is motivated by its resemblance to the spatial-frequency response characteristic of retinal ganglion cells [103, 30], and also helps gradient based learning by uniforming variances along the main axes of variation. A statistically independent ‘validation’ set of 64,000 whitened natural images was created in the same way and used to monitor and report validation losses during model training (Fig. 2a). Another statistically independent ‘test’ set of 64,000 whitened natural images was also created in an identical way and was used throughout the paper for model evaluation.

### Texture image datasets

Our texture image datasets, created solely for model evaluation, were synthesized from 15 texture family seed images with the algorithm in [40]. Computational constraints limited the training image patch sizes to 40×40 or 50×50 pixels; therefore, we selected only starting images for which the dominant wavelength of synthesized textures fits into a 40×40 patch. We then randomly cropped a balanced set of 40×40-pixel or 50×50-pixel texture patches from the synthesized texture images, matched the grand total pixel histogram to the standard normal distribution for each texture family separately, and applied the whitening procedure used for natural images to the whole texture dataset. Fig. 2c shows sample patches (before whitening) from the selected 15 texture families. Using this procedure, we created a balanced ‘train’ texture dataset of 600,000 patches in total and a statistically independent balanced ‘test’ texture dataset of 60,000 patches in total and used these for the texture family decoding experiments in Figs. 2d and 6d. In the rest of the paper, only the latter, ‘test’ texture dataset was used for model evaluation. (For more details, see [23].) For our CORnet discriminative benchmark we used non-whitened 224×224-pixel patches created in a similar way.

### Phase scrambling

Phase scrambling (Figs. 3g and 6f) was performed on the cropped 40×40 images (either from the van Hateren dataset or the synthesized texture image dataset) in the way described in [16] (and previously in [40]): we took the Fourier transform of the image, randomized the phase of each Fourier component, and took the real part of the inverse Fourier transform of the result. This manipulation did not change the spectral properties of the input images. After this we applied histogram matching and ZCA whitening as with the non-phase scrambled images.

### Filter scrambling

Filter scrambling (Fig. 6f) was performed in a way that is equivalent to low-level-synthetic (LL-synthetic) image generation in [47], to which it is compared in Fig. 6g. First, we calculated the Z_1_ posterior mean of the image under the shallow-VAE model, randomly permuted unit activations separately for active and non-active units, and generated the mean image from the result with the shallow-VAE model. This manipulation did not preserve the spectral properties of the input images. Finally, histogram matching and ZCA whitening were applied as above.

### Comparison texture image dataset

Figs. 2e, 3c, 3e, and 3g compare TDVAE model predictions to experiments in [16] and [17]. These two papers are built on the same electrophysiology data set that we tried to mimic as described here. The papers present spike counts in a 100 ms time window for 102 individual V1 and 103 individual V2 neurons found in 13 macaque monkeys as a response to the same texture stimulus sequence with 15 texture families, 15 samples plus 15 phase scrambled samples from each texture family, and 20 presentations per image. In our comparison dataset, we randomly sampled 100 active Z_1_ dimensions as a model of the 102 V1 neurons in the experiment. The TDVAE model only has 6 active dimensions in Z_2_ which is less than the 103 experimental V2 neurons. As a substitute, we generated 100 random rotations of the 6D space and took the first components of the rotated 6D active Z_2_ space as a model of 100 V2 neurons. (Note that this procedure is consistent with the rotation-invariant unit normal prior in the Z_2_ layer of the TDVAE model.) To avoid correlated responses which are clearly not present in the experimental dataset, we sampled the stimulus-conditional model posteriors separately for all 100 Z_1_ and 100 Z_2_ model neurons, respectively. For stimuli, we used test texture images from the 15 texture families used throughout this paper, 15 samples plus 15 phase scrambled samples from each texture family, and 20 presentations per image. We named this dataset the ‘comparison’ dataset. To mimic the 100 ms time window in the experiment, we sampled the model posteriors in the following way. For Z_1_ units, we took one sample from the Z_2_ posterior and used that single Z_2_ sample in the Z_1_ posterior from which we averaged five samples. For Z_2_ units, we simply took a sample from the Z_2_ posterior.

### Linear and non-linear characterization of model units

*Receptive fields* of Z_1_ and Z_2_ units (Figs. 2b, 4a, 5d, 6c, Extended Data Figs. 6, 7, 8) were calculated with a method customary in visual neuroscience, reverse correlation estimated by spike-triggered averaging (STA, [104]), using 128,000 white noise images where each individual pixel intensity value was drawn from an independent Gaussian distribution with its position and scale matching the grand total mean and standard deviation of the pixels in the ‘test’ natural image dataset. In the case of the CORnet model, receptive fields of 64 independent features (corresponding to the depth of the convolutional map) were calculated. The receptive fields of the remaining features are translated versions of the computed ones due to the convolutional nature of the latent space. *Projective fields* (Extended Data Figs. 6, 8) were calculated with latent traversal along each model unit in Z_1_ and Z_2_ by taking two points symmetrically to 0 that differed only along the given unit dimension, generating a large number (Z_1_: 128, Z_2_: 2560) of mean images from these two points and taking the difference between the mean images for the two points. The *second-order receptive fields* of a Z_2_ unit (Fig. 2b) were calculated with the spike-triggered covariance (STC) method as described in [104]. We plot two representative eigenvectors of the STC matrix. We had to use 10^8^ white noise images to get visible structure.

### Identifying active model units

In a given model, we classified Z_1_ and Z_2_ model units into active and inactive categories based on four criteria (Extended Data Fig. 6). (1) Variance of the mean response to the images in the ‘test’ natural image dataset was found to be orders of magnitude larger for active units than for inactive units. (2) Mean of the response variance (uncertainty) over the same image dataset also creates two clean clusters for active and inactive units. (3) Mean squared pixel intensities of receptive fields are clearly larger for active units than for inactive units. (4) Mean squared pixel intensities of projective fields are also larger for active units than for inactive units. We found that these four criteria consistently cluster units into the same active and inactive sets (for TDVAE, see Extended Data Fig. 6). The number of active Z_1_ and Z_2_ units is denoted in Extended Data Table 4 for all discussed models. In the paper we only consider active model units.

### Equivalence of Z_1_ receptive fields in single-layer and hierarchical models

Learned Z_1_ receptive fields are localized, oriented and bandpass in both the hierarchical TDVAE model and the single-layer shallow-VAE model (Extended Data Figs. 7, 8). We calculated two quantities for each active Z_1_ unit for a more detailed comparison: filter center (estimated as the center of mass of the pointwise squared receptive field) and dominant wave vector (calculated from the maximum position of the squared magnitude of the Fourier transform of the receptive field). To increase the precision of these estimates, we used projective fields instead of receptive fields as they are very similar but less noisy (Extended Data Fig. 8). We found that the empirical distribution of filter centers and dominant wave vectors are very similar in the TDVAE and shallow-VAE models, making them equivalent complete bases in the space of natural training images, at least in terms of first-order probability densities.

### Multiunit texture family decoding

Linear decoding accuracies of texture family in Figs. 2d, 5a, 5c, and 6d were calculated with multinomial logistic regression. Each bar reports the mean of test classification accuracies calculated from *n* = 5 fitting sessions started from different random seeds. Standard deviation values are not plotted since they are all below 0.01 and would be invisible. In each case, the dependent variable was texture family and independent variables were derived from the ‘train’ and ‘test’ texture datasets as follows. Fig. 2d: *Pixels*: independent variables were image pixel intensities. Z_1_ *and* Z_2_ *of all VAE models*: independent variables were the mean responses of all active units in the denoted layer of the denoted model to ‘train’ and ‘test’ texture images; for the untrained model, we used the same number of randomly selected units. *CORnet-*Z_1_: independent variables were CORnet-Z V1 ‘conv’ layer units in response to non-whitened 224×224-pixel texture images. To make comparison with our generative models, only model units which correspond to the central 40×40-pixel part of the image were considered (20×20×64 units). *CORnet-*Z_2_: independent variables were 10×10×128 CORnet-Z V2 ‘conv’ layer units. Figs. 5a and 5c: mean responses of all active Z_1_ units in TDVAE. Fig. 6d: Z_1_ *means:* mean responses of 100 randomly selected active Z_1_ units. Z_2_ *means:* mean responses of the six active units. Z_1_ *corrs* (*original textures, phase scrambled, filter scrambled*): independent variables were noise correlation coefficients between the mean responses of the same 100 randomly selected active Z_1_ units as for Z_1_ means, calculated from 16,384 presentations of every 10th image in the original, phase scrambled and filter scrambled texture image datasets, respectively. Overfitting was prevented with weight decay regularization where needed.

### Robust texture representation in the Z_2_ model layer

The multiunit texture family decoding test was repeated for mean Z_1_ and Z_2_ responses in all models presented in the paper. As it is shown in Extended Data Table 5, decoding accuracies from Z_1_ were around chance level 1/15 = 0.067 for all models (we have 15 texture families) and decoding accuracies from Z_2_ were close to perfect performance 1 for all hierarchical models, implying no texture representation in Z_1_ and a robust texture representation in Z_2_ for all studied models. Repeating the same analysis for subsets of active Z_2_ units showed that texture family is only decodable from some of the Z_2_ dimensions and not from others in TDVAE models (last two columns of Extended Data Table 5), revealing a low-dimensional (4–7 dimensions) learned texture representation in TDVAE models. In contrast, we found no axis-alignment in the Z_2_ texture representation in the ffVAE model as texture family could be more or less decoded from all active Z_2_ units (Extended Data Table 5). Since our primary model, TDVAE with deep nonlinearity, contains no non-texture-family-coding active Z_2_ units, we do not analyze such units in the present paper. We also found that even in the discriminative model CORnet, decoding accuracy from the V1 layer was just above chance and decoding accuracy from the V2 layer was high.

### t-SNE visualization

In Fig. 2e we compare the low dimensional visualization of model responses in the TDVAE model with the low dimensional visualization of neural responses in [17] (Fig. 2f). To this end, we used the ‘comparison’ dataset defined above and the t-SNE visualization method utilized in [17]. t-distributed stochastic neighbor embedding (t-SNE) [105] attempts to minimize the divergence between the distributions of neighbor probability in the original high dimensional data space and the visualized low dimensional space. Similar to [17], we input a set of 225 data vectors (15 texture families, 15 images per family) to the t-SNE algorithm, each of which collected the mean responses of 100 neurons in a model layer to a stimulus. Like [17], we normalized the data so that, for each model unit, responses to the 225 images had a mean of 0 and standard deviation of 1, and executed the t-SNE analysis with a perplexity value of 30.

### ANOVA

We also repeated the ANOVA study in [17] in the context of the TDVAE model (Figs. 3c and 3d). The utilized nested ANOVA method partitioned the total variance of Z_1_ and Z_2_ model unit responses to the ‘comparison’ dataset into portions arising across families, across samples within a family, and across repetitions of the same stimulus. The ANOVA generates an F-statistic that captures the ratio of variances between each hierarchical level. For all model neurons we found the F-statistic to be significant for ratios of variance across repetitions and across samples as well as for ratios of variance across samples and across families. Following [17] we performed further analysis using the ratio between partitioned variance components. To obtain the variance ratio, we divided the percent variance across families by the percent variance across samples.

### Classification decoding

We also reproduced the decoding study in [17] with the TDVAE model (Figs. 3e and 3f; differences from [17] in parentheses). We used Gaussian decoders (experiment: Poisson decoders) to classify model responses to the ‘comparison’ dataset into 15 sample categories per family and, independently, into 15 texture family categories. On each iteration, we randomly selected a number of units from our model neurons in the Z_1_ or Z_2_ model layer. To compute performance in the sample classification task, we estimated the mean and variance of the response of each selected neuron for each of the 15 samples within each family by computing the sample mean and sample variance over four randomly selected repetitions out of the 20 repetitions in the dataset (experiment used 10 repetitions but that made the task trivial for the model). For the held-out 16 repetitions of each sample, we computed which of the 15 samples was most likely to have produced the population response, assuming independent normal distributions for the model units under the estimated mean and variance. We computed the average performance (% correct) over all samples and families, and repeated this process 100 times to get a performance for each population size (experiment used 10,000 here but estimated errors for the model were already small at 100 repetitions). To compute performance in the family classification task, we estimated the mean and variance of the response for each family over 8 randomly selected samples from the 15 different samples and for all repetitions. For each of the repetitions of the held-out seven samples, we computed which of the five families was most likely to have produced the population response. We computed the average performance over all repetitions and repeated this process 100 times to get a performance for each population size (experiment used 10,000 here but estimated errors for the model were already small at 100 repetitions). We computed performance measures for both tasks using population sizes of 1, 3, 10, 30, and 100 neurons.

### Phase scrambling induced modulation of response magnitudes in Z_1_ and Z_2_

We studied how reducing high-level stimulus structure affects model responses by reproducing the electrophysiology experiment in [16] (see Figs. 3g and 3h). First, as the experiment queries changes in response magnitudes, we took the absolute value of all model responses to undisturbed and phase scrambled texture images in the ‘comparison’ dataset. Following [16], we then calculated per-neuron per-family mean response magnitudes by averaging these absolute responses over per-family samples and stimulus presentations. Next we calculated modulation indices for each ‘comparison’ Z_1_ and Z_2_ unit and texture family by dividing the difference between the above mean response magnitudes to undisturbed and phase scrambled textures with their sum. As the final step, we averaged these per-neuron modulation indices over the 15 texture families and plotted separate histograms for Z_1_ and Z_2_ units (Fig. 3g).

### Filter selection for top-down experiments

While studying top-down effects with the TDVAE model, we found that the results were sensitive to certain limitations of the model such as pixel aliasing and finite patch size. To get rid of the ensuing artifacts, we restricted our top-down experiments to Z_1_ dimensions with central, localized, medium wavelength generative fields, which we selected algorithmically with the following criteria: 1) centrality: center of mass of the pointwise square of the generative field falls in the central (40%)^2^> of the image patch, 2) locality: size *σ* of the filter, calculated by fitting a cylindrical Gaussian profile with scale *σ* to the pointwise absolute value of the Z_1_ generative field by requiring the two to have the same moment of inertia, falls between 18% and 33% of the patch edge length, 3) medium-wavelength: the dominant wavelength, calculated from the maximum position of the squared magnitude of the 2D Fourier transform of the generative field, falls between 17% and 28% of the patch edge length. These criteria found 40 (40×40-pixel TDVAE), 57 (50×50-pixel TDVAE), and 53 (50×50-pixel shallowVAE) central, localized, medium wavelength Z_1_ filters, respectively, which we used exclusively in Figs. 4, 5, and 6 as well as Extended Data Figs. 9b,c,d and 10, except for the texture family decoding experiments in Figs. 5a,c and Fig. 6d. For the discriminative CORnet model we started out from the 64 Z_1_-like units that had their 7×7-pixel receptive fields in the center of the 224×224-pixel input images. For these, all illusory-like stimuli fitted into the 224×224-pixel input image area. However, the receptive fields of some of these 64 Z_1_-like units were not localized and oriented, preventing the alignment of the illusory stimuli to them. To algorithmically select only localized and oriented filters, we leveraged a key mathematical property of Gabor wavelets: their uncertainty, defined as the product of their variances in the position and wave number domains, reaches the theoretical minimum of 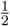.Based on this criterion, we selected the 20 units with the lowest-uncertainty receptive fields.

### Illusory contour stimuli

Stimuli used for illusory contour experiments (Figs. 4a-g and 5b, Extended Data Fig. 9b) were based on stimuli used in the experimental paper [43] (see Fig. 4c) that were aligned to the 57 (50×50-pixel TDVAE) or 53 (50×50-pixel shallowVAE) central, localized, medium wavelength Z_1_ filters (see above). Upright and upside-down stimuli were both used, hence the *n* = 114 Z_1_ units in Fig. 4d. We applied a second, principled filter selection step that left only Z_1_ units with significantly amplified illusory TDVAE model responses compared to the linear response. The selection criteria were the following: 1) fit into patch: the line segment connecting the two “Pac-Man” centers in the ‘Illusory’ stimulus aligned to the receptive field (based on the above definitions of center of mass and dominant wave vector), extended by 11.1% in both directions, fits into the patch, 2) overlap between the ‘Illusory’ stimulus and the generative field (defined as the sum of the squared generative field pixel values on the figure positions of the ‘Illusory’ stimulus divided by the same sum but masked with the ‘Line’ stimulus instead) is less than 0.1, 3) all four response peaks in Fig. 4e are closer than 0.25 pixels to each other (this is a proxy of regular peak shapes). This selection step found *n* = 9 Z_1_ units for 50×50-pixel TDVAE, *n* = 29 Z_1_ units for 50×50-pixel shallowVAE, and *n* = 0 Z_1_ units for 40×40-pixel TDVAE. In technical terms, generative-field-fitted ‘Line’, ‘Illusory’, and ‘Rotated’ stimulus sets were first generated in black (−1) and white (1) with 16×16 grid supersampling, then downsampled in a bilinear way, filtered with Gaussian blur, Z scored, and whitened with the PCA basis obtained from natural image whitening above. (Whitening with the PCA basis of natural images instead of the illusory contour PCA basis was chosen because of the small effective dimensionality of the illusory contour stimulus set.) Illusory stimulus parameters used in Figs. 4 and 5b: Pac-Man radius: 9 pixels; Gaussian blur standard deviation: 0.9 pixels; step size of stimulus displacement: 0.125 pixels. For the discriminative CORnet model we used the following stimulus parameters: Pac-Man radius: 3 pixels; Gaussian blur standard deviation: 1.0 pixels; step size of stimulus displacement: 0.5 pixels. Note that, since CORnet was trained on raw, not-whitened images, we did not apply whitening to the illusory stimuli in this case.

### Model responses to illusory stimuli

One important goal of modeling illusory contour responses in Figs. 4 and 5b was to faithfully compare them with Fig. 3a in the experimental paper [43] (reproduced as Fig. 4c). This Figure in [43] presents mean spike counts in a 50 ms time window, a response averaging procedure we modeled in the following way. Experimentally, autocorrelations of membrane potentials for static stimuli typically decay on a ~20 ms timescale in the V1 cortical area [34]. Therefore, in the context of Figs. 4 and 5b, we modeled 50 ms response averaging in [43] by averaging three independent samples from Z_1_ model response distributions. Autocorrelation times in higher cortical areas are longer [106, 107, 108, 109], which we modeled by taking the three Z_1_ samples using the very same sample from Z_2_ model response distributions. For consistency, all model responses to illusory stimuli in Figs. 4 and 5b, as well as Extended Data Fig. 9 were calculated in this way. Ref. [43] does not specify the number of trials each stimulus was presented to the animal. We performed 500 trials for each illusory stimulus in our model experiments which proved high enough that most selected model units showed a significant illusory effect (boosting or suppression). On the other hand, CORnet is a deterministic model, therefore, we could only calculate one model response value for each stimulus–model unit pair. This means that while we were able to calculate differences between model and linear responses, we could not make statements on the significance of their differences.

### Contour completion stimuli

Stimuli used for the contour completion experiment (Fig. 5d, Extended Data Figs. 9c,d) were based on the stimuli used in the experimental paper [45] with the exception that the lattice constant (distance between neighboring bar centers) was reduced by 10% to help the stimuli fit into a 50×50-pixel patch. As in [45], the central bars of the stimuli were aligned to the 57 central, localized, medium wavelength Z_1_ filters found in the 50×50-pixel TDVAE model (see above for the filter selection procedure). Like the illusory contour stimuli, contour completion stimuli were first generated in black (−1) and white (1) with 16×16 grid supersampling, then downsampled in a bilinear way, filtered with Gaussian blur, Z scored, and whitened with the PCA basis obtained from natural image whitening above. (Whitening with the PCA basis of natural images instead of the contour completion PCA basis was chosen because of the small effective dimensionality of the contour completion stimulus set.) Contour completion stimulus parameters used in Fig. 5d were the following: bar length: 11 pixels; Gaussian blur standard deviation: 0.9 pixels. Extended Data Fig. 9c uses a bar length of 13 pixels instead of 11 pixels, and Extended Data Fig. 9d a Gaussian blur standard deviation of 1 pixel instead of 0.9 pixels. A contrast value (defined as the standard deviation of pixel intensities in the whitened stimuli) of 0.8 was used everywhere.

### Model responses to contour completion stimuli and their analysis

The goal of our contour completion experiment was to prove the existence of top-down boosting in the responses of central, localized, medium wavelength Z_1_ units of the 50×50-pixel TDVAE model to filter-aligned contour completion stimuli. Therefore, we used an analysis similar to our illusory contour experiment above, as follows. For each Z_1_ filter, each stimulus type (‘random segments’ and ‘three parallel segments’, see Fig. 5d), and each stimulus parameter set (bar length, Gaussian sigma, contrast) we generated 100 stimuli with different random orientations of the freely rotating bars. We selected an experimental time window of 50 ms and used the same one-Z_2_-sample mean-of-three-Z_1_-samples model sampling procedure as with the illusory contour experiment. Using this procedure, we conducted *N* = 30 trials for each stimulus (like in [45]). For each stimulus and Z_1_ unit, we normalized the trial-averaged three-parallel-segments model response with the trial-averaged random-orientations model response, and subtracted from it the three-parallel-segments linear response normalized with the random-orientations linear response. As the final step, we selected the values of bar length, Gaussian blur standard deviation, and contrast, and for each Z_1_ unit we plotted the mean of these differences over the 100 random stimulus orientations described above, each Z_1_ unit giving one dot in Fig. 5d, and Extended Data Fig. 9c,d, respectively.

### Noise and signal correlations

We calculated noise correlation matrices by recording the mean responses of the 40 central, localized, medium wavelength Z_1_ filters in the 40×40-pixel TDVAE model to the repeated presentation of the same stimulus and then calculating the Pearson product-moment correlation coefficients between each distinct pair of Z_1_ units over stimulus presentations. We calculated the signal correlation matrix of a texture family by selecting a number of samples from the family, recording the means of the mean responses of the 40 central, localized, medium wavelength Z_1_ filters in the 40×40-pixel TDVAE model to the repeated presentation of each sample, and calculating the Pearson product-moment correlation coefficients between each distinct pair of Z_1_ units over the selected samples. Following [47], we defined the dissimilarity of two correlation matrices as the mean absolute difference between their elements. To mimic the 400 ms recording window used in [47], we repeated the above experiments with three more model sampling procedures: mean of 20 Z_1_ samples from one Z_2_ sample; mean of 2×10 Z_1_ samples; and mean of 4×5 Z_1_ samples (Extended Data Fig. 10c,d). On a final note, we only used natural images for calculating correlations that had a contrast (standard deviation of pixel intensity values) larger than the observation noise value 0.4 used when training our VAE models.

### Statistics

To test the significance of our results, we used one- and two-sample, one- and twosided, independent- and paired-sample versions of Student’s t-test, depending on the specific individual questions we were asking about the data. We documented the details of the individual statistical tests together with their results throughout the paper.

## Acknowledgements

The authors thank Mihály Bányai and Dávid G. Nagy for early discussions about the model. G.O. was supported by a grant from the Human Frontiers Science Program, and by the European Union project RRF-2.3.1-21-2022-00004 within the framework of the Artificial Intelligence National Laboratory in Hungary and the National Research, Development and Innovation Office under grant agreement ADVANCED 150361. We thank the HUN-REN Cloud for providing compute support for the study (see [110]; https://science-cloud.hu/). We are grateful for RyeongKyung Yoon and Xaq Pitkow and other three anonymous reviewers for their highly constructive comments to the manuscript.

## Author contributions

F.C., B.M, and G.O. conceptualized the study, F.C. performed data curation, F.C. and B.M. wrote simulation software, F.C. performed the simulations, F.C., B.M., and K.Ó. performed the analyses, F.C. and G.O. were designing the visualization, F.C., G.O., and B.M. wrote the manuscript.

## Competing interests

The authors declare no competing interests.

**Extended Data Fig. 1.**
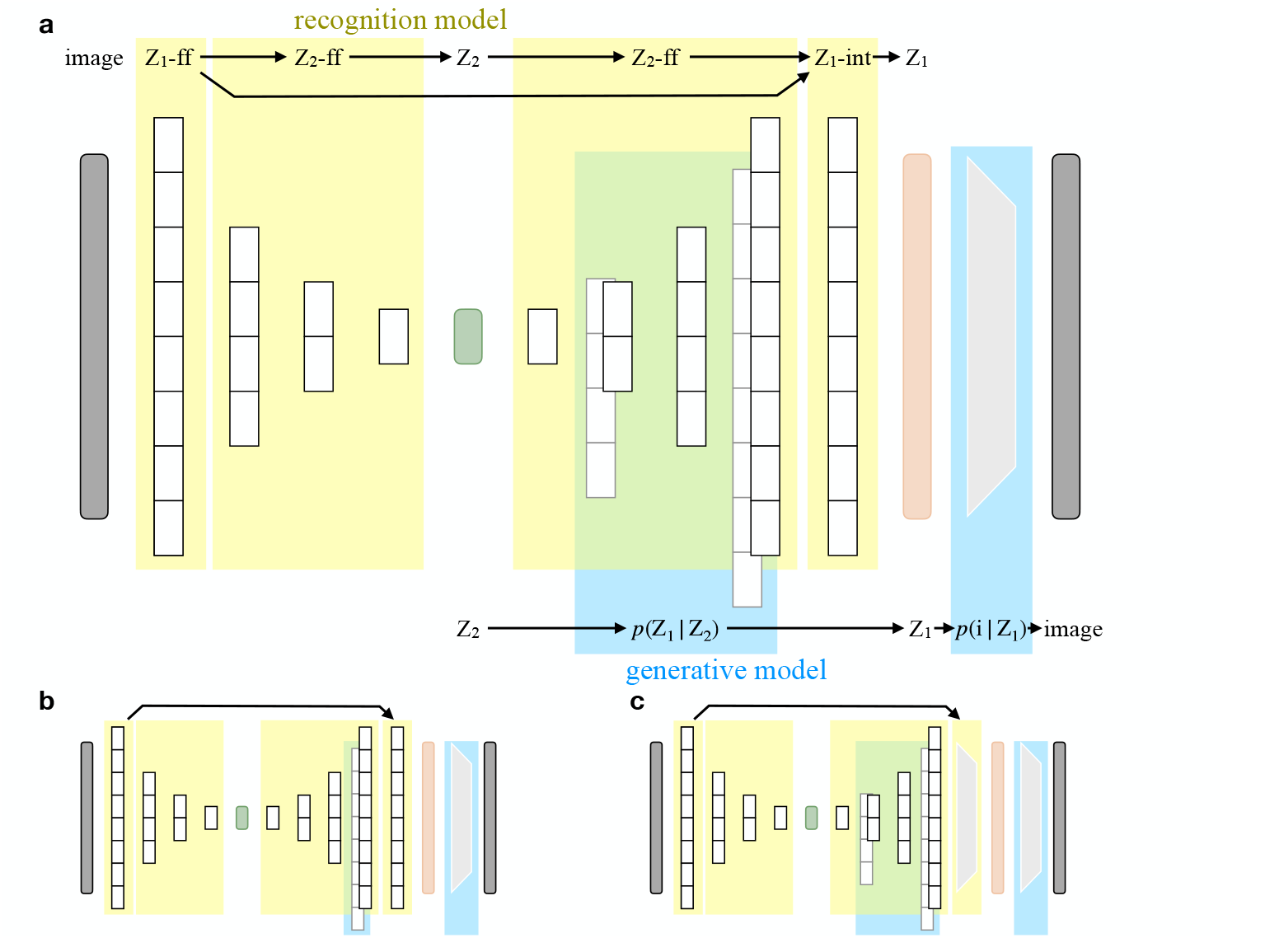
Computational graphs of TDVAE model variants. **a**, Computational architecture for our main model (TDVAE with deep nonlinearity). Each shaded object that includes multiple hidden layers represents an MLP which transforms its input in a nonlinear manner. Grey trapezoids represent linear transformations. Yellow corresponds to the recognition part of the model, blue corresponds to the generative model. **b**, Computational graph of TDVAE with shallow nonlinearity. **c**, Computational architecture of TDVAE with linear integration.

**Extended Data Table 2.** Architecture of models with bottom-up recognition. Number of hidden units in each MLP layer computing the mean and the standard deviation of each conditional distribution in the bottom-up recognition models.

| Model name | $p_{\theta}(Z_1 \vert Z_2)$ | $q_{\Phi}(Z_1 \vert \text{image})$ | $q_{\Phi}(Z_2 \vert Z_1)$ |
| --- | --- | --- | --- |
| shallow-VAE | N/A | (2000, 2000) | N/A |
| ffVAE | (2000) | (2000, 2000) | (1000, 500, 250) |
| shallow-VAE-50px | N/A | (3000, 3000) | N/A |

**Extended Data Table 3.** Architecture of models with top-down recognition. Number of hidden units in each MLP layer computing the mean and the standard deviation of each conditional distribution in TDVAE models. In the non-shared model, the MLP Z_2_-ff takes its input directly from pixels, eliminating parameter sharing in the entire computation graph.

| Model name | $p_\theta(Z_1 \vert Z_2)$ | $Z_1$-ff | $Z_2$-ff | $Z_1$-TD | $Z_1$-INT |
| --- | --- | --- | --- | --- | --- |
| TDVAE<br>linear integration | (1000, 2000) | (2000) | (1000, 500, 250) | (250, 500, 1000, 2000) | () |
| TDVAE<br>shallow nonlinearity | (2000) | (2000) | (1000, 500, 250) | (250, 500, 1000, 2000) | (2000) |
| TDVAE<br>deep nonlinearity | (1000, 2000) | (2000) | (1000, 500, 250) | (250, 500, 1000, 2000) | (2000) |
| TDVAE-50px<br>shallow nonlinearity | (3000) | (3000) | (1500, 750, 500) | (500, 750, 1500, 3000) | (3000) |
| TDVAE-20px | (250, 500) | (500) | (250, 125, 63) | (63, 125, 250, 500) | (500) |
| TDVAE-20px<br>non-shared | (250, 500) | (500) | (500, 250, 125, 63) | (63, 125, 250, 500) | (500) |

**Extended Data Table 4.** Basic features of the trained models. Validation ELBO at the end of training and the number of active stochastic units in the trained Z_1_ and Z_2_ model layers for each presented model.

| Model name | ELBO | active $Z_1$ units | active $Z_2$ units |
| --- | --- | --- | --- |
| shallow-VAE | −1916 | 1256 | N/A |
| TDVAE linear integration | −1886 | 1256 | 27 |
| ffVAE shallow nonlinearity | −1818 | 1256 | 25 |
| TDVAE shallow nonlinearity | −1705 | 1256 | 199 |
| TDVAE deep nonlinearity | −1699 | 1256 | 6 |
| TDVAE deep nonlinearity untrained | −17,319 | N/A | N/A |
| shallow-VAE-50px | −3111 | 1963 | N/A |
| TDVAE-50px shallow nonlinearity | −2753 | 1963 | 7 |
| TDVAE-20px | −400 | 314 | 4 |
| TDVAE-20px non-shared | −407 | 314 | 2 |

**Extended Data Table 5.**
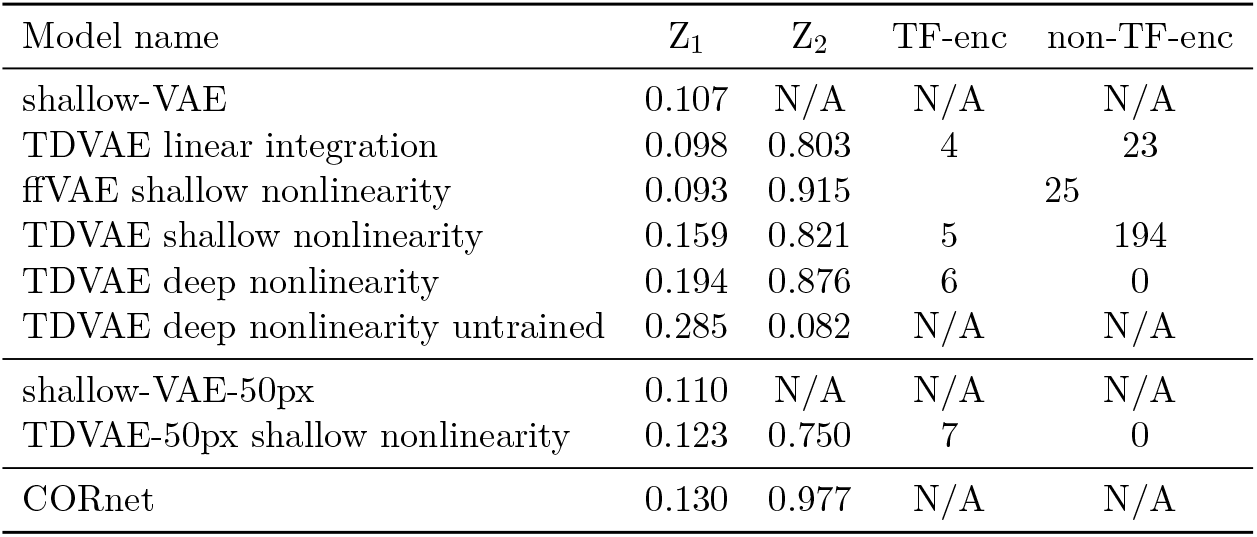
Texture representation in the trained models. Texture family decoding accuracies from mean responses of active Z_1_ and active Z_2_ units and the number of texture-family-coding (TF-enc) and non-texture-family-coding (non-TF-enc) active Z_2_ units for all studied models. No clear distinction was found between texture-family-coding and non-texture-family-coding Z_2_ units in the ffVAE model.

**Extended Data Fig. 6.**
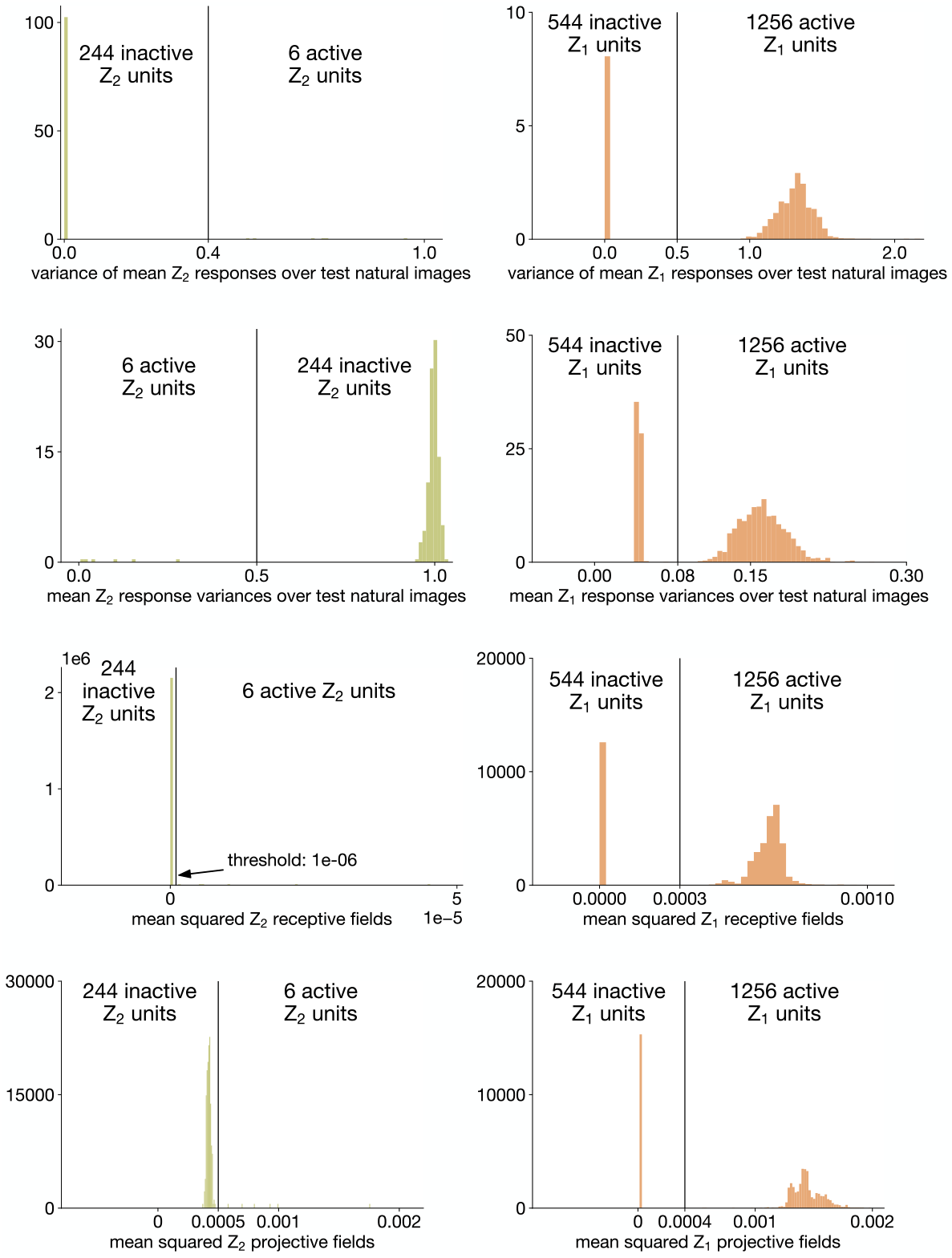
Activity of model units in the trained TDVAE model. We used four criteria for selecting active model units in the Z_1_ and Z_2_ model layers (Methods): (from top to bottom) stimulus dependent responses (variance of the mean response over the ‘test’ natural image dataset); response confidence (mean of the response variance over the ‘test’ natural image dataset); response to white noise (mean of the squared receptive field); influence on generated images (mean of the squared projective field). We found that all criteria consistently selected the same six active units in Z_2_ (left) and that, in a similar way, all criteria consistently selected the same 1256 active units in Z_1_ (right). Note that the ‘training’ natural image dataset has 1256 effective dimensions (Methods), meaning that the Z_1_ layer learned a complete linear basis from and over the training data.

**Extended Data Fig. 7.**
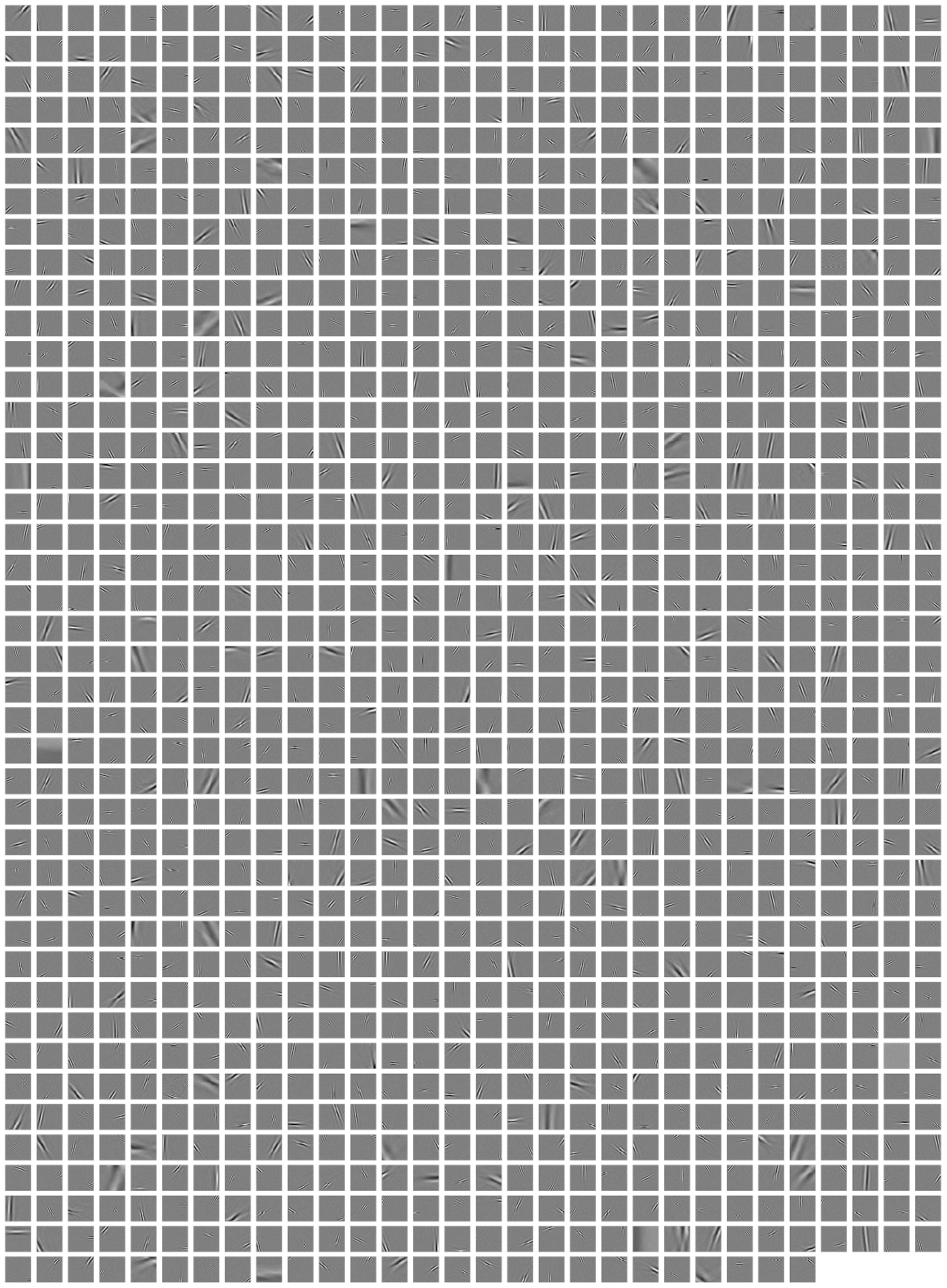
Receptive fields of all active Z_1_ units in the TDVAE model. All Z_1_ receptive fields are localized, oriented, and bandpass, reminiscent of V1 simple cells in primates.

**Extended Data Fig. 8.**
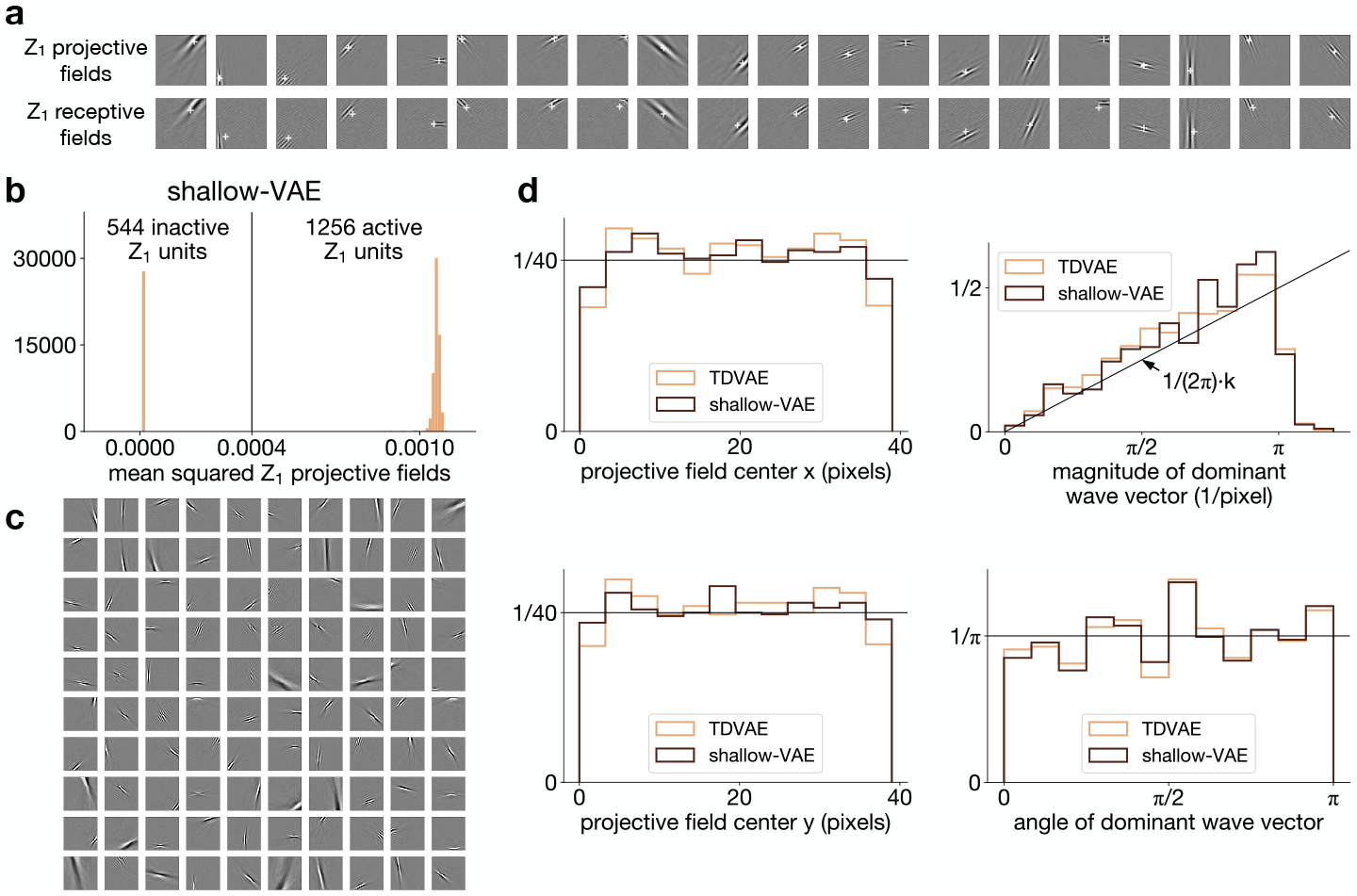
Equivalence of Z_1_ receptive fields in the TDVAE and the shallow-VAE models. **a**, Example Z_1_ projective and receptive field centers calculated in the TDVAE model with the same procedure (Methods). Projective fields are very similar to receptive fields but also smoother, making them more reliable for center of mass estimation than receptive fields. In general, we use projective fields for estimating Z_1_ filter parameters throughout the paper. **b**, The shallow-VAE model has 1256 active Z_1_ units, like the TDVAE model. **c**, Example projective fields from the shallow-VAE model. These are localized, oriented, and bandpass, like those in the TDVAE model. **d**, Z_1_ projective field centers and dominant wave vectors (Methods) homogeneously fill the patch area in the TDVAE and the shallow-VAE models, making them equivalent complete bases in the space of natural training images, at least in terms of first-order probability densities.

**Extended Data Fig. 9.**
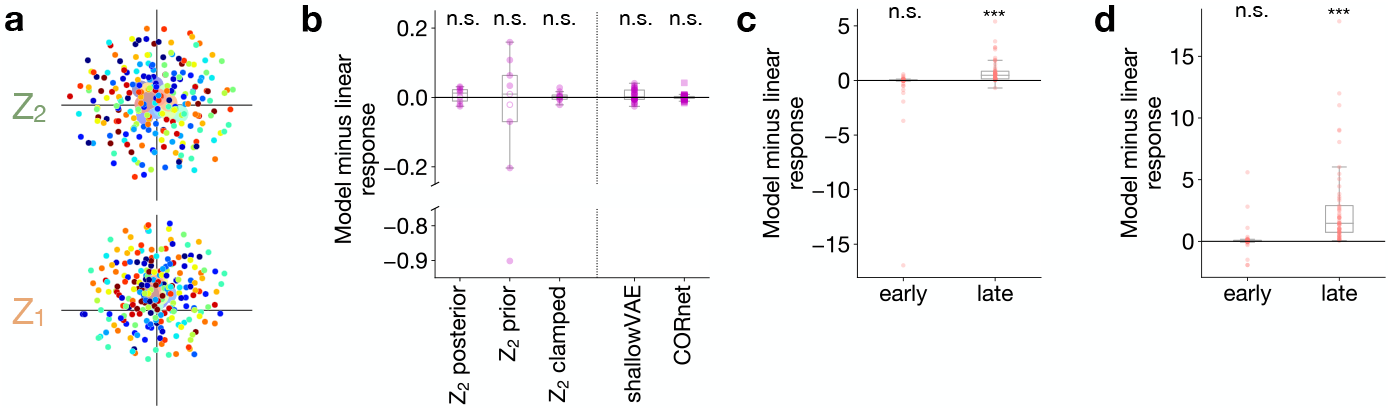
Representation and top-down effects in the TDVAE model. **a**, Two-dimensional visualization (t-SNE) of mean responses of Z_1_ and Z_2_ neurons to the texture images used in Fig. 2e in the untrained TDVAE model (*dots*). *Colors* indicate the 15 texture families as on panels Fig. 2c and 2e. *Disks*: mean across different samples from the same family. **b**, Differences between the mean and linear responses to the ‘Rotated’ stimulus (center line, median; box limits, upper and lower quartiles; whiskers, 1.5× interquartile range). For notation, see Fig. 4g. All t-tests are two-sided, one-sample against 0. Z_2_ *posterior*: *n* = 9; mean: 0.0065, P = 0.39, t = 0.92, df = 8. Z_2_ *prior*: *n* = 9; mean: −0.091, P = 0.42, t = −0.85, df = 8. Z_2_ *clamped*: *n* = 9; mean: 0.0028, P = 0.56, t = 0.60, df = 8. *shallowVAE*: *n* = 29; mean: 0.0061, P = 0.070, t = 1.9, df = 28. *CORnet*: *n* = 40; mean: 9.7 × 10^−5^, P = 0.94, t = 0.070, df = 39. **c**, The contour completion experiment in Fig. 5d repeated with longer bars (bar lengths were 13 pixels instead of 11 pixels; center line, median; box limits, upper and lower quartiles; whiskers, 1.5× interquartile range). Only the late response shows significant boosting (*n* = 57 in both cases; early: mean effect size: −0.40, two-sided one-sample t-test against 0: P = 0.19, t = −1.3, df = 56; late: mean effect size: 0.77, one-sided one-sample t-test against 0: P = 4.2× 10^−7^, t = 5.5, df = 56). **d**, The contour completion experiment in Fig. 5d repeated with a larger Gaussian kernel (kernel size was 1 pixel instead of 0.9 pixels; center line, median; box limits, upper and lower quartiles; whiskers, 1.5× interquartile range). Only the late response shows significant boosting (*n* = 57 in both cases; early: mean effect size: 0.11, two-sided one-sample t-test against 0: P = 0.40, t = 0.86, df = 56; late: mean effect size: 2.7, one-sided one-sample t-test against 0: P = 9.1 × 10^−8^, t = 6.0, df = 56).

**Extended Data Fig. 10.**
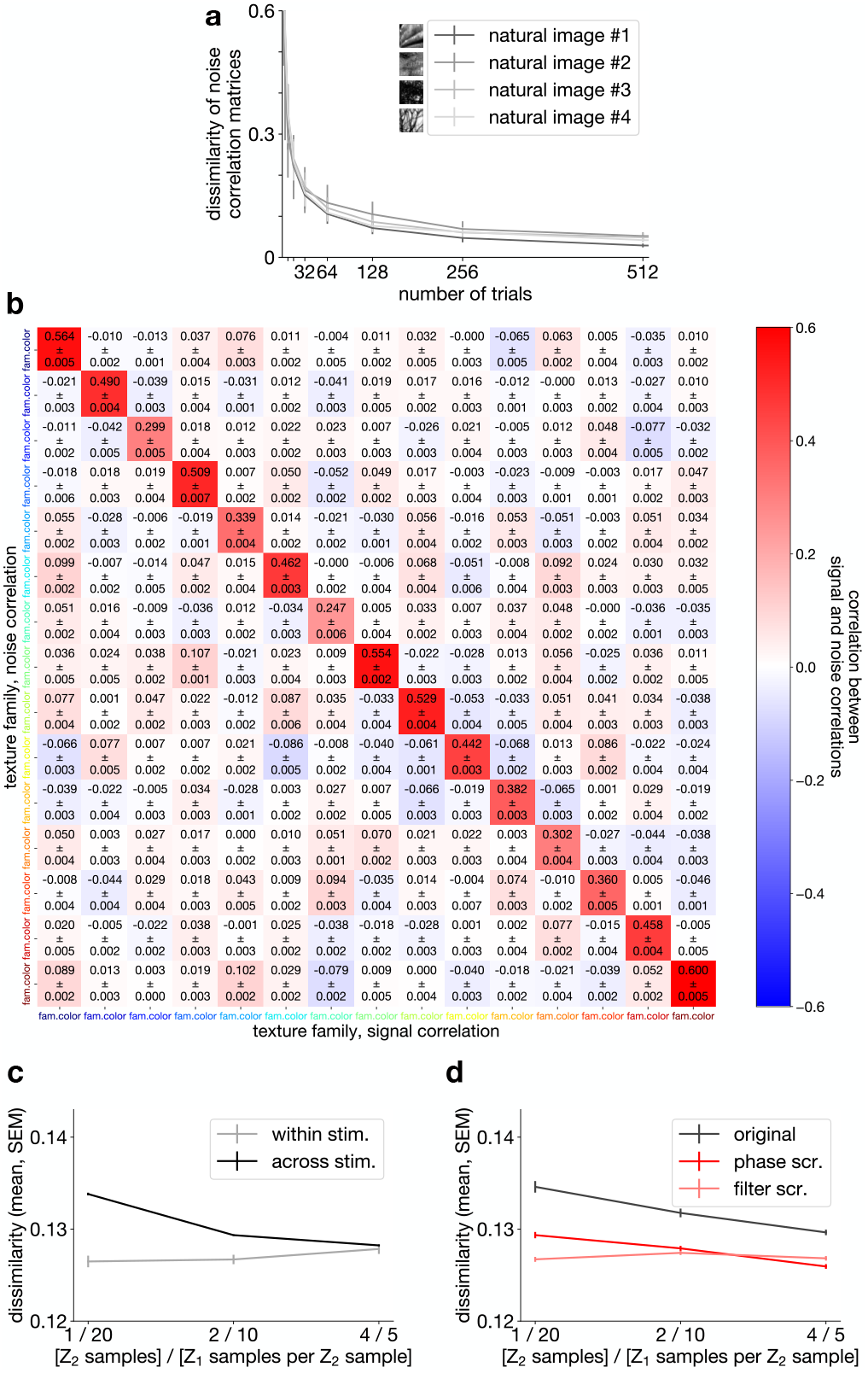
Noise and signal correlations in the TDVAE model. **a**, Estimation of noise correlations improves with the number of trials they are calculated from. Solid lines: mean within-stimulus noise correlation matrix dissimilarities for four natural image examples (*inset*). Error bars: s.d. In general, this paper presents noise correlation matrices calculated from *n* = 80 trials. **b**, Noise and signal correlations are connected through the contextual prior. Pearson correlations between noise and signal correlation coefficients over distinct pairs of *n* = 40 Z_1_ units. Correlation matrices were calculated on 15 texture families (axes), 80 samples per family, and 80 trials per sample. The whole process was repeated 5 times. Colors: means. Errors: s.e.m. **c**, Fig. 6b repeated with different model sampling settings compatible with the 400 ms recording window used in [47]. **d**, Fig. 6f repeated with different model sampling settings compatible with the 400 ms recording window used in [47].

## References

[1] Richards, B. A. et al. A deep learning framework for neuroscience. Nature Neuroscience 22, 1761–1770 (2019).

[2] Yamins, D. L. & DiCarlo, J. J. Using goal-driven deep learning models to understand sensory cortex. Nature Neuroscience 19, 356–365 (2016).

[3] Bashivan, P., Kar, K. & DiCarlo, J. J. Neural population control via deep image synthesis. Science 364, eaav9436 (2019).

[4] Walker, E. Y. et al. Inception loops discover what excites neurons most using deep predictive models. Nature Neuroscience 22, 2060–2065 (2019).

[5] Kar, K., Kubilius, J., Schmidt, K., Issa, E. B. & DiCarlo, J. J. Evidence that recurrent circuits are critical to the ventral stream’s execution of core object recognition behavior. Nature Neuroscience 22, 974–983 (2019).

[6] Conwell, C. et al. Neural regression, representational similarity, model zoology & neural taskonomy at scale in rodent visual cortex. Advances in Neural Information Processing Systems 34, 5590–5607 (2021).

[7] Lange, Richard D., Shivkumar, Sabyasachi, Chattoraj, Ankani, & Haefner Ralf M. Bayesian encoding and decoding as distinct perspectives on neural coding. Nature Neuroscience 26(12), 2063–2072 (2023).

[8] De Lange, F. P., Heilbron, M. & Kok, P. How do expectations shape perception? Trends in Cognitive Sciences 22, 764–779 (2018).

[9] Kersten, D., Mamassian, P. & Yuille, A. Object perception as Bayesian inference. Annu. Rev. Psychol. 55, 271–304 (2004).

[10] Lee, T. S. & Mumford, D. Hierarchical Bayesian inference in the visual cortex. JOSA A 20, 1434–1448 (2003).

[11] Körding, K. P. & Wolpert, D. M. Bayesian integration in sensorimotor learning. Nature 427, 244–247 (2004).

[12] Berkes, P., Orbán, G., Lengyel, M. & Fiser, J. Spontaneous cortical activity reveals hallmarks of an optimal internal model of the environment. Science 331, 83–87 (2011).

[13] Gibson, James J. The ecological approach to visual perception. Moughton Mifflin (1979).

[14] Julesz, Bela Textons, the elements of texture perception, and their interactions. Nature 290(5802), 91–97 (1981).

[15] Freeman, J. & Simoncelli, E. P. Metamers of the ventral stream. Nature Neuroscience 14, 1195–1201 (2011).

[16] Freeman, J., Ziemba, C. M., Heeger, D. J., Simoncelli, E. P. & Movshon, J. A. A functional and perceptual signature of the second visual area in primates. Nature Neuroscience 16, 974–981 (2013).

[17] Ziemba, C. M., Freeman, J., Movshon, J. A. & Simoncelli, E. P. Selectivity and tolerance for visual texture in macaque V2. Proceedings of the National Academy of Sciences 113, E3140–E3149 (2016).

[18] Okazawa, Gouki, Tajima, Satohiro, & Komatsu, Hidehiko Image statistics underlying natural texture selectivity of neurons in macaque V4. Proceedings of the National Academy of Sciences 112(4), E351–E360 (2015).

[19] Bolaños, F. et al. Efficient coding of natural images in the mouse visual cortex. bioRxiv (2022). URL https://www.biorxiv.org/content/early/2022/09/17/2022.09.14.507893.

[20] Khaligh-Razavi, S.-M. & Kriegeskorte, N. Deep supervised, but not unsupervised, models may explain IT cortical representation. PLoS Computational Biology 10, e1003915 (2014).

[21] Nayebi, A. et al. Mouse visual cortex as a limited resource system that self-learns an ecologically-general representation. PLOS Computational Biology 19, e1011506 (2023).

[22] Zhuang, C. et al. Unsupervised neural network models of the ventral visual stream. Proceedings of the National Academy of Sciences 118, e2014196118 (2021).

[23] Csikor, F., Meszéna, B., Szabó, B. & Orban, G. Top-down effects in an early visual cortex inspired hierarchical Variational Autoencoder. In SVRHM 2022 Workshop @ NeurIPS (2022). URL https://openreview.net/forum?id=8dfboOQfYt3.

[24] Gilbert, C. D. & Li, W. Top-down influences on visual processing. Nature Reviews Neuroscience 14, 350–363 (2013).

[25] Kingma, D. P. & Welling, M. Auto-encoding Variational Bayes (2013). URL 10.48550/arXiv.1312.6114.

[26] Rezende, D. J., Mohamed, S. & Wierstra, D. Stochastic backpropagation and approximate inference in deep generative models. In Xing, E.P. & Jebara, T. (eds.) Proceedings of the 31st International Conference on Machine Learning, vol. 32 of Proceedings of Machine Learning Research, 1278–1286 (PMLR, Bejing, China, 2014). URL https://proceedings.mlr.press/v32/rezende14.html.

[27] Child, R. Very deep VAEs generalize autoregressive models and can outperform them on images (2021). URL 10.48550/arXiv.2011.10650.

[28] Vahdat, A. & Kautz, J. NVAE: A deep hierarchical variational autoencoder. Advances in Neural Information Processing Systems 33, 19667–19679 (2020).

[29] Olshausen, B. A. & Field, D. J. Emergence of simple-cell receptive field properties by learning a sparse code for natural images. Nature 381, 607–609 (1996).

[30] Geadah, Victor, Barello, Gabriel, Greenidge, Daniel, Charles Adam S, & Pillow Jonathan W Sparse-coding variational autoencoders. Neural Computation 36, 2571–2601 (2024).

[31] Knill, D. C. & Richards, W. Perception as Bayesian inference (Cambridge University Press, 1996).

[32] Yuille, A. & Kersten, D. Vision as Bayesian inference: analysis by synthesis? Trends in Cognitive Sciences 10, 301–308 (2006).

[33] Fiser József, Berkes, Pietro Orbán, Gergő, & Lengyel, Máté Statistically optimal perception and learning: from behavior to neural representations. Trends in Cognitive Sciences 14, 119–130 (2010).

[34] Azouz, R. & Gray, C. M. Cellular mechanisms contributing to response variability of cortical neurons in vivo. Journal of Neuroscience 19, 2209–2223 (1999).

[35] Cohen, Marlene R, & Kohn, Adam Measuring and interpreting neuronal correlations. Nature Neuroscience 14, 811–819 (2011).

[36] Orbán, G., Berkes, P., Fiser, J. & Lengyel, M. Neural variability and sampling-based probabilistic representations in the visual cortex. Neuron 92, 530–543 (2016).

[37] Festa, D., Aschner, A., Davila, A., Kohn, A. & Coen-Cagli, R. Neuronal variability reflects probabilistic inference tuned to natural image statistics. Nature Communications 12, 1–11 (2021).

[38] Carandini, M. Amplification of trial-to-trial response variability by neurons in visual cortex. PLoS Biology 2, e264 (2004).

[39] Yu, Y., Stirman, J. N., Dorsett, C. R. & Smith, S. L. Selective representations of texture and motion in mouse higher visual areas. Current Biology 32, 2810–2820 (2022).

[40] Portilla, J. & Simoncelli, E. P. A parametric texture model based on joint statistics of complex wavelet coefficients. International Journal of Computer Vision 40, 49–70 (2000).

[41] Kubilius, J. et al. Cornet: Modeling the neural mechanisms of core object recognition. BioRxiv 408385 (2018). URL https://www.biorxiv.org/content/10.1101/408385v1.

[42] Weiss, Y., Simoncelli, E. P. & Adelson, E. H. Motion illusions as optimal percepts. Nature Neuroscience 5, 598–604 (2002).

[43] Lee, T. S. & Nguyen, M. Dynamics of subjective contour formation in the early visual cortex. Proceedings of the National Academy of Sciences 98, 1907–1911 (2001).

[44] Ziemba, Corey M, Perez Richard K, Pai, Julia, Kelly Jenna G, Hallum Luke E, Shooner, Christopher, & Movshon, J Anthony Laminar differences in responses to naturalistic texture in macaque V1 and V2. Journal of Neuroscience 39, 9748–9756 (2019).

[45] Chen, Minggui, Yan, Yin, Gong, Xiajing, Gilbert Charles D., Liang, Hualou, & Li, Wu Incremental integration of global contours through interplay between visual cortical areas. Neuron 82(3), 682–694 (2014).

[46] Ringach, Dario L Spontaneous and driven cortical activity: implications for computation. Current Opinion in Neurobiology 19, 439–444 (2009).

[47] Bányai, M. et al. Stimulus complexity shapes response correlations in primary visual cortex. Proceedings of the National Academy of Sciences 116, 2723–2732 (2019).

[48] Karklin, Y. & Lewicki, M. S. Emergence of complex cell properties by learning to generalize in natural scenes. Nature 457, 83–86 (2009).

[49] Coen-Cagli, R., Kohn, A. & Schwartz, O. Flexible gating of contextual influences in natural vision. Nature Neuroscience 18, 1648–1655 (2015).

[50] Fu, J. et al. Pattern completion and disruption characterize contextual modulation in mouse visual cortex. bioRxiv (2023). URL https://www.biorxiv.org/content/early/2023/03/14/2023.03.13.532473.

[51] Boutin, Victor, Franciosini, Angelo, Ruffier, Franck, & Perrinet, Laurent Effect of top-down connections in Hierarchical Sparse Coding. Neural Computation 32(11), 2279–2309 (2020).

[52] Boutin, Victor, Franciosini, Angelo, Chavane, Frederic, Ruffier, Franck, & Perrinet, Laurent Sparse deep predictive coding captures contour integration capabilities of the early visual system. PLoS Computational Biology 17(1), e1008629 (2021).

[53] Schwartz, O. & Simoncelli, E. P. Natural signal statistics and sensory gain control. Nature Neuroscience 4, 819–825 (2001).

[54] Karklin, Yan, & Lewicki Michael S Learning higher-order structures in natural images. Network: Computation in Neural Systems 14, 483–499 (2003).

[55] Vafaii, Hadi, Galor, Dekel, & Yates Jacob L. Poisson Variational Autoencoder. arXiv preprint 2405.14473 (2024). URL https://www.arxiv.org/abs/2405.14473.

[56] Higgins, I. et al. Unsupervised deep learning identifies semantic disentanglement in single inferotemporal face patch neurons. Nature Communications 12, 1–14 (2021).

[57] Vafaii, Hadi, Yates, Jacob, & Butts, Daniel Hierarchical VAEs provide a normative account of motion processing in the primate brain. Advances in Neural Information Processing Systems 36, (2024).

[58] Burgess, C. P. et al. Understanding disentangling in β-VAE (2018). URL 10.48550/arXiv.1804.03599.

[59] Hoyer, P. & Hyvärinen, A. Interpreting neural response variability as Monte Carlo sampling of the posterior. Advances in Neural Information Processing Systems 15 (2002).

[60] Haefner, R. M., Berkes, P. & Fiser, J. Perceptual decision-making as probabilistic inference by neural sampling. Neuron 90, 649–660 (2016).

[61] Shrinivasan, Suhas, Lurz, Konstantin-Klemens, Restivo, Kelli, Denfield, George, Tolias, Andreas, Walker, Edgar, & Sinz, Fabian Taking the neural sampling code very seriously: A data-driven approach for evaluating generative models of the visual system. Advances in Neural Information Processing Systems 36 (2024).

[62] Bányai, M. & Orbán, G. Noise correlations and perceptual inference. Current Opinion in Neurobiology 58, 209–217 (2019).

[63] Echeveste, R., Aitchison, L., Hennequin, G. & Lengyel, M. Cortical-like dynamics in recurrent circuits optimized for sampling-based probabilistic inference. Nature Neuroscience 23, 1138–1149 (2020).

[64] Hennequin, G., Ahmadian, Y., Rubin, D. B., Lengyel, M. & Miller, K. D. The dynamical regime of sensory cortex: stable dynamics around a single stimulus-tuned attractor account for patterns of noise variability. Neuron 98, 846–860 (2018).

[65] Soo, Wayne, & Lengyel, Máté Training stochastic stabilized supralinear networks by dynamics-neutral growth. Advances in Neural Information Processing Systems 35, 29278– 29291 (2022).

[66] Hosoya, Haruo, & Hyvärinen, Aapo A hierarchical statistical model of natural images explains tuning properties in V2. Journal of Neuroscience 35, 10412–10428 (2015).

[67] Catoni, Josefina, Martos, Domonkos, Csikor, Ferenc, Ferrante, Enzo, Milone Diego H, Meszéna, Balázs, Orbán, Gergő, & Echeveste, Rodrigo Uncertainty in latent representations of variational autoencoders optimized for visual tasks. arXiv preprint 2404.15390 (2025). URL https://www.arxiv.org/abs/2404.15390.

[68] Rao, R. P. & Ballard, D. H. Predictive coding in the visual cortex: a functional interpretation of some extra-classical receptive-field effects. Nature Neuroscience 2, 79–87 (1999).

[69] Lotter, W., Kreiman, G. & Cox, D. A neural network trained for prediction mimics diverse features of biological neurons and perception. Nature Machine Intelligence 2, 210–219 (2020).

[70] Ali, A., Ahmad, N., de Groot, E., van Gerven, M. A. J. & Kietzmann, T. C. Predictive coding is a consequence of energy efficiency in recurrent neural networks. Patterns 3 (2022).

[71] Walker, E. Y., Cotton, R. J., Ma, W. J. & Tolias, A. S. A neural basis of probabilistic computation in visual cortex. Nature Neuroscience 23, 122–129 (2020).

[72] Ernst, M. O. & Banks, M. S. Humans integrate visual and haptic information in a statistically optimal fashion. Nature 415, 429–433 (2002).

[73] Boutin, V., Zerroug, A., Jung, M. & Serre, T. Iterative VAE as a predictive brain model for out-of-distribution generalization. In NeurIPS 2020 Workshop SVRHM (2020). URL https://openreview.net/forum?id=jE6SlVTOFPV.

[74] Aitchison, L. & Lengyel, M. With or without you: predictive coding and Bayesian inference in the brain. Current Opinion in Neurobiology 46, 219–227 (2017).

[75] Markov, Nikola T, Vezoli, Julien, Chameau, Pascal, Falchier, Arnaud, Quilodran, René, Huissoud, Cyril, Lamy, Camille, Misery, Pierre, Giroud, Pascale, & Ullman, Shimon, et al. Anatomy of hierarchy: feedforward and feedback pathways in macaque visual cortex. Journal of Comparative Neurology 522, 225–259 (2014).

[76] Keller, Andreas J., Roth Morgane M., & Scanziani, Massimo Feedback generates a second receptive field in neurons of the visual cortex. Nature 582(7813), 545–549 (2020).

[77] Kirchberger, Lisa, Mukherjee, Sreedeep, Self Matthew W., & Roelfsema Pieter R. Contextual drive of neuronal responses in mouse V1 in the absence of feedforward input. Science Advances 9(3), eadd2498 (2023).

[78] Kubilius, Jonas, Schrimpf, Martin, Kar, Kohitij, Rajalingham, Rishi, Hong, Ha, Majaj, Najib, Issa, Elias, Bashivan, Pouya, Prescott-Roy, Jonathan, Schmidt, Kailyn, & others Brain-like object recognition with high-performing shallow recurrent ANNs. Advances in Neural Information Processing Systems 32, (2019).

[79] Rajaei, Karim, Mohsenzadeh, Yalda, Ebrahimpour, Reza, & Khaligh-Razavi, Seyed-Mahdi Beyond core object recognition: Recurrent processes account for object recognition under occlusion. PLoS Computational Biology 15(5), e1007001 (2019).

[80] Osindero, Simon, & Hinton Geoffrey E. Modeling image patches with a directed hierarchy of Markov random fields. Advances in Neural Information Processing Systems 20 (2007).

[81] Wei, Xue-Xin, & Stocker Alan A. Efficient coding provides a direct link between prior and likelihood in perceptual Bayesian inference. Advances in Neural Information Processing Systems 25 (2012).

[82] Ganguli, Deep, & Simoncelli Eero P. Efficient sensory encoding and Bayesian inference with heterogeneous neural populations. Neural Computation 26(10), 2103–2134 (2014).

[83] Vacher, Jonathan, Launay, Claire, & Coen-Cagli, Ruben Flexibly regularized mixture models and application to image segmentation. Neural Networks 149, 107–123 (2022).

[84] Lange, R. D. & Haefner, R. M. Task-induced neural covariability as a signature of approximate Bayesian learning and inference. PLoS Computational Biology 18, e1009557 (2022).

[85] Harel, A., Kravitz, D. J. & Baker, C. I. Task context impacts visual object processing differentially across the cortex. Proceedings of the National Academy of Sciences 111, E962–E971 (2014).

[86] Lange, R. D. & Haefner, R. M. Characterizing and interpreting the influence of internal variables on sensory activity. Current Opinion in Neurobiology 46, 84–89 (2017).

[87] Bondy, A. G., Haefner, R. M. & Cumming, B. G. Feedback determines the structure of correlated variability in primary visual cortex. Nature Neuroscience 21, 598–606 (2018).

[88] Lazar, A. et al. Paying attention to natural scenes in area V1. bioRxiv (2023). URL https://www.biorxiv.org/content/early/2023/08/26/2023.03.21.533636.

[89] Baker, Nicholas, Erlikhman, Gennady, Kellman Philip J., & Lu, Hongjing Deep Convolutional Networks do not Perceive Illusory Contours. In CogSci (2018).

[90] Baker, Nicholas, Lu, Hongjing, Erlikhman, Gennady, & Kellman Philip J. Deep convolutional networks do not classify based on global object shape. PLoS Computational Biology 14(12), e1006613 (2018).

[91] Pang, Zhaoyang, O’May Callum Biggs, Choksi, Bhavin, & VanRullen, Rufin Predictive coding feedback results in perceived illusory contours in a recurrent neural network. Neural Networks 144, 164–175 (2021).

[92] Kogo, Naoki & Froyen, Vicky Emergence of border-ownership by large-scale consistency and long-range interactions: Neuro-computational model to reflect global configurations. Psychological Review 128(4), 597 (2021).

[93] Knebel, Jean-François & Murray Micah M. Towards a resolution of conflicting models of illusory contour processing in humans. NeuroImage 59(3), 2808–2817 (2012).

[94] Pak, Alexandr, Ryu, Esther, Li, Claudia, & Chubykin Alexander A. Top-down feedback controls the cortical representation of illusory contours in mouse primary visual cortex. Journal of Neuroscience 40(3), 648–660 (2020).

[95] George, Dileep & Hawkins, Jeff Towards a mathematical theory of cortical micro-circuits. PLoS Computational Biology 5(10), e1000532 (2009).

[96] Dura-Bernal, Salvador, Wennekers, Thomas, & Denham Susan L. Top-down feedback in an HMAX-like cortical model of object perception based on hierarchical Bayesian networks and belief propagation. PLOS ONE 7(11), e48216 (2012).

[97] Młynarski, W., Hledík, M., Sokolowski, T. R. & Tkačik, G. Statistical analysis and optimality of neural systems. Neuron 109, 1227–1241 (2021).

[98] Pillow, Jonathan & Rubin, Nava Perceptual completion across the vertical meridian and the role of early visual cortex. Neuron 33(5), 805–813 (2002).

## References

[99] Kingma, D. P. & Welling, M. An introduction to Variational Autoencoders. Foundations and Trends® in Machine Learning 12, 307–392 (2019). URL 10.1561/2200000056.

[100] Gáspár Merse E., Polack, Pierre-Olivier, Golshani, Peyman, Lengyel, Máté, & Orbán, Gergő Representational untangling by the firing rate nonlinearity in V1 simple cells. Elife 8, e43625 (2019).

[101] Pascanu, R., Mikolov, T. & Bengio, Y. On the difficulty of training recurrent neural networks. In Dasgupta, S. & McAllester, D. (eds.) Proceedings of the 30th International Conference on Machine Learning, vol. 28 of Proceedings of Machine Learning Research, 1310– 1318 (PMLR, Atlanta, Georgia, USA, 2013). URL https://proceedings.mlr.press/v28/pascanu13.html.

[102] Van Hateren, J. H. & van der Schaaf, A. Independent component filters of natural images compared with simple cells in primary visual cortex. Proceedings of the Royal Society of London. Series B: Biological Sciences 265, 359–366 (1998).

[103] Olshausen, B. A. & Field, D. J. Sparse coding with an overcomplete basis set: A strategy employed by V1? Vision Research 37, 3311–3325 (1997). URL https://www.sciencedirect.com/science/article/pii/S0042698997001697.

[104] Rust, N. C., Schwartz, O., Movshon, J. A., Simoncelli, E. P., Spatiotemporal elements of macaque V1 receptive fields. Neuron 46, 945–956 (2005).

[105] van der Maaten, L. & Hinton, G. Visualizing data using t-SNE. Journal of Machine Learning Research 9 (2008).

[106] Murray, J. D. et al. A hierarchy of intrinsic timescales across primate cortex. Nature Neuroscience 17, 1661–1663 (2014).

[107] Runyan, C. A., Piasini, E., Panzeri, S. & Harvey, C. D. Distinct timescales of population coding across cortex. Nature 548, 92–96 (2017).

[108] Siegle, J. H. et al. Survey of spiking in the mouse visual system reveals functional hierarchy. Nature 592, 86–92 (2021).

[109] Brunec, I. K. & Momennejad, I. Predictive representations in hippocampal and prefrontal hierarchies. Journal of Neuroscience 42, 299–312 (2022).

[110] Héder, M. et al. The Past, Present and Future of the ELKH Cloud, Információs Társadalom 22, 128 (2022).

